# Minimal functional domains of the core polarity regulator Dlg

**DOI:** 10.1101/2022.04.29.490107

**Authors:** Mark J. Khoury, David Bilder

## Abstract

The compartmentalized domains of polarized epithelial cells arise from mutually antagonistic actions between the apical Par complex and the basolateral Scrib module. In *Drosophila*, the Scrib module proteins Scribble (Scrib) and Discs-large (Dlg) are required to limit Lgl phosphorylation at the basolateral cortex, but how Scrib and Dlg could carry out such a ‘protection’ activity is not clear. We tested Protein Phosphatase 1α (PP1) as a potential mediator of this activity but demonstrate that a significant component of Scrib and Dlg regulation of Lgl is PP1-independent and found no evidence for a Scrib-Dlg-PP1 protein complex. However, the Dlg SH3 domain plays a role in Lgl protection and, in combination with the N-terminal region of the Dlg HOOK domain, in recruitment of Scrib to the membrane. We identify a ‘minimal Dlg’ comprised of the SH3 and HOOK domains that is both necessary and sufficient for Scrib localization and epithelial polarity function *in vivo*.

**Summary Statement:** A minimal SH3-HOOK fragment of Dlg is sufficient to support epithelial polarity through mechanisms independent of the PP1 phosphatase.

## INTRODUCTION

Cell polarity is the fundamental process by which a single cell partitions its plasma membrane into two molecularly distinct, mutually exclusive domains. The ability to polarize is crucial for the development and homeostasis of many cell types, including neurons, stem cells and epithelial cells (St Johnston and Ahringer, 2010). Epithelial cells exhibit apicobasal polarity, a feature critical for their physiological function and morphogenesis of their resident tissues (Buckley and St Johnston, 2022; Rodriguez-Boulan and Macara, 2014). Like many other polarized cells, epithelial cell polarity is often regulated by two highly conserved groups of proteins: the Par complex, composed of Par-3, Par-6 and atypical protein kinase C (aPKC) and the Scrib module, composed of Scribble (Scrib), Discs-large (Dlg) and Lethal giant larvae (Lgl) (Flores-Benitez and Knust, 2016; Goldstein and Macara, 2007). The separation of apical and basolateral domains derives from the mutual antagonism between the apical-defining Par complex and the basolateral-defining Scrib module. Apical aPKC phosphorylates Lgl, which removes it from the plasma membrane, thus excluding Lgl from the apical domain (Bailey and Prehoda, 2015; Betschinger et al., 2003; Dong et al., 2015; Plant et al., 2003). Conversely, basolateral Lgl inhibits aPKC localization to prevent apical domain spread (Hutterer et al., 2004; Wirtz-Peitz et al., 2008; Yamanaka et al., 2003).

For the Par complex, there is now detailed insight into specific functions and molecular interactions for each of its component proteins (Lang and Munro, 2017; Tepass, 2012). By contrast, how the Scrib module determines basolateral polarity is poorly defined (Bonello and Peifer, 2018; Nakajima, 2021; Stephens et al., 2018). The major knowledge gap in Scrib module biology is the molecular mechanism of Scrib and Dlg activity. While Lgl’s role as an antagonist of aPKC localization is well-known, how Scrib and Dlg act to ensure restriction of the apical domain is not understood. Addressing this question will be essential to a full understanding of cell polarity. We previously identified several principles of Scrib module protein function, showing that

Dlg is required to regulate Scrib cortical localization and providing evidence that Scrib and Dlg are both required to negatively regulate Lgl phosphorylation (Khoury and Bilder, 2020). The data led us to propose a model in which Scrib and Dlg act as molecular switches in the aPKC-Lgl relationship. At the basolateral domain, Scrib and Dlg ‘protect’ Lgl by limiting inhibitory aPKC phosphorylation, allowing Lgl to antagonize aPKC, whereas at the apical domain, where Scrib and Dlg are not present, Lgl is unprotected and can be inhibited by aPKC. Here, we have used a combination of *in vivo* genetics, biochemistry, and an *in vitro* polarity system to pursue potential molecular bases of this model. We fail to find evidence supporting a plausible mechanism of Lgl protection in which Scrib and Dlg recruit the phosphatase PP1, but we identify a minimal domain of Dlg that is both necessary and sufficient for Scrib recruitment and polarity function.

## RESULTS

### PP1 is a candidate effector of Scrib and Dlg activity

Since Scrib and Dlg are both scaffolding proteins, it is likely that any Lgl protection activity derives from specific binding partners. To search for polarity-relevant Dlg binding partners, we have carried out APEX2-based proximity proteomics of Dlg in epithelial tissue (Sharp et al., 2021). We mined these data for potential effectors of Lgl protection and uncovered the *Drosophila* Protein Phosphatase 1α (PP1α) homolog, Pp1-87B (hereafter PP1), in the top 60 most enriched Dlg-proximity hits (log2 fold change=4.8, p=0.001). PP1 is an appealing candidate to mediate Lgl regulation by Scrib and Dlg because PP1 was recently shown to counteract aPKC phosphorylation of Lgl in *Drosophila* epithelial cells (Moreira et al., 2019). We confirmed that *pp1* depletion resulted in decreased cortical Lgl localization in follicle epithelial cells (**Fig. S1C-E**). Intriguingly, both Scrib and Dlg contain conserved protein sequences that match PP1-binding consensus motifs: Scrib contains SILK and RVxF motifs and Dlg contains an RVxF motif (**Fig. 1A, Fig. S2A**) (Hendrickx et al., 2009; Young et al., 2013). These motifs are found in proteins that bind PP1 and can act as substrate specificity factors, recruiting the general PP1 phosphatase to target proteins (Heroes et al., 2013). We therefore hypothesized that Scrib and Dlg could regulate Lgl phosphorylation by scaffolding PP1 at the basolateral cortex.

**Figure 1.**
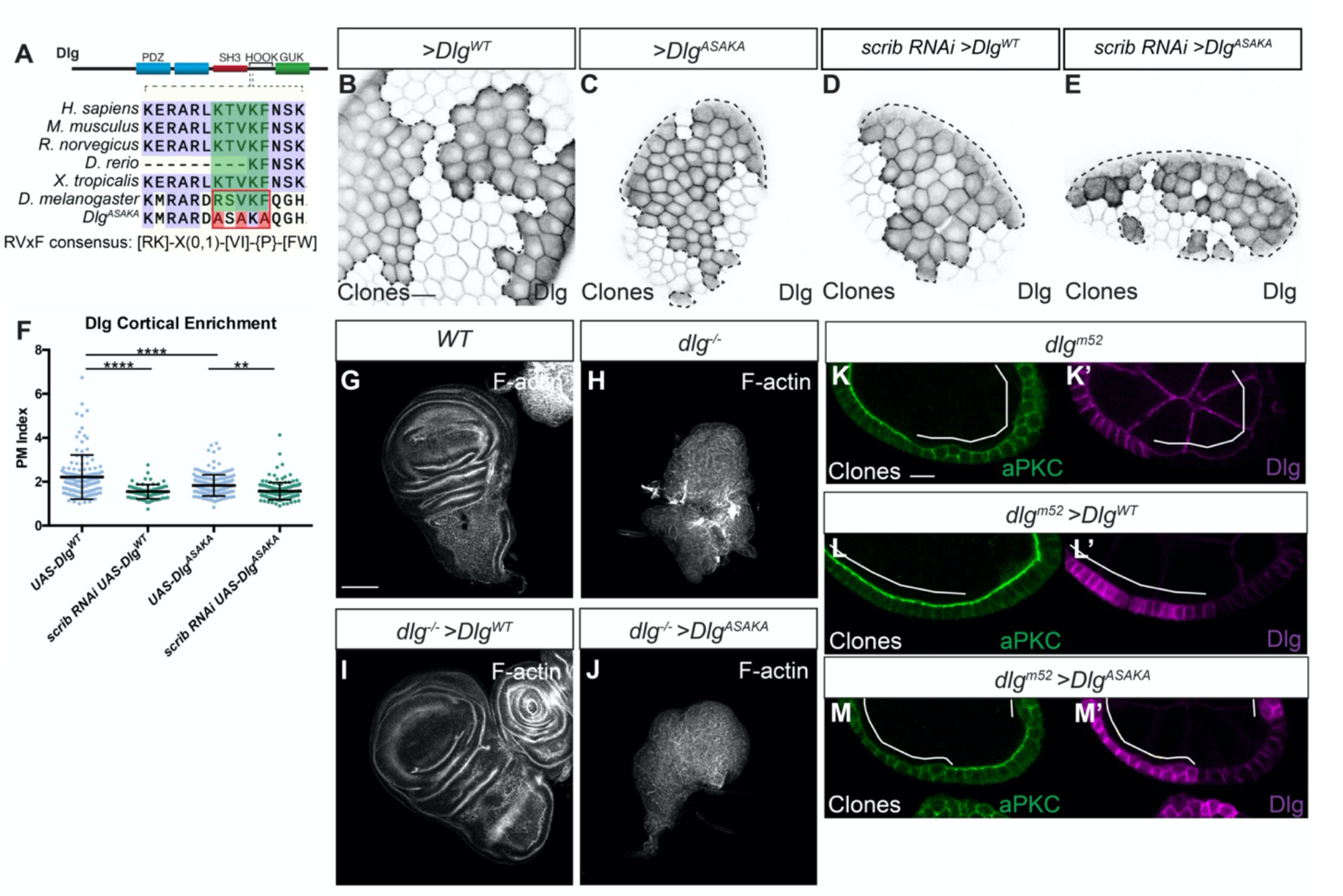
The Dlg RVxF motif is critical for function. (A) Cartoon showing location of the RVxF motif in the Dlg protein and conservation of the motif across species. The resides mutated in Dlg^ASAKA^ are highlighted in red. The RVxF consensus as defined by (Wakula et al., 2003) is shown. (B-E) Like Dlg^WT^ (B), Dlg^ASAKA^ localizes to the basolateral membrane and is enriched at the cell cortex (C). Dlg^WT^ localization (D) as well as Dlg^ASAKA^ localization (E) is sensitive to *scrib*-depletion, quantified in (F). (G-J) Compared to WT (G), *dlg* null mutant wing discs form disorganized tumors (H). Expression of Dlg^WT^ rescues this phenotype (I), while expression of Dlg^ASAKA^ does not rescue (J). (K-M) In the follicle epithelium, *dlg*^*m52*^ null mutants (K) lose polarity, characterized by lateral aPKC spread, and this is rescued by expressing Dlg^WT^ (L). In contrast, polarity loss is not rescued by Dlg^ASAKA^ expression (M). Scale bars, 10μm except in G-J, 100μm. Dotted lines in (B-E) and white lines in (K-M) indicate clones of given genotype. (F) One-way ANOVA with Tukey’s multiple comparisons test. Error bars represent S.D. Data points are PM Index measurements in single cells. PM Index=cortical/cytoplasmic intensity. **P < 0.01, ****P < 0.0001.

### Functional tests of Scrib and Dlg PP1-binding motifs

To test the functional relevance of the putative PP1-binding motifs in Scrib and Dlg, we designed targeted mutations in these sequences. We first mutated the consensus residues of the Dlg RVxF motif to alanine (**Fig. 1A**). This construct (Dlg^ASAKA^) localized to the basolateral membrane in follicle cells and was enriched at the cell cortex (PM Index > 1) (**Fig. 1B-C,F**). Dlg^ASAKA^ localization was slightly less cortical than overexpressed WT Dlg (**Fig. 1F**). However, Dlg^ASAKA^ localization was still sensitive to *scrib* depletion, suggesting that this mutation does not prevent the recently described electrostatic mechanism of Dlg membrane recruitment (**Fig. 1D-F**) (Lu et al., 2021). Dlg^ASAKA^ did not rescue the overproliferation or polarity defects in *dlg* mutant wing imaginal discs (**Fig. 1G-J**). Similarly, when expressed in *dlg* mutant follicle cell clones, Dlg^ASAKA^ had no rescuing activity and these cells were indistinguishable from *dlg* null mutants, with ectopic basolateral aPKC localization and epithelial multilayering (**Fig. 1K-M**). Thus, the Dlg RVxF motif is required for all tested Dlg functions.

Next, we mutated the critical residues in the Scrib SILK and RVxF motifs to alanine (**Fig. S2A**). As the SILK motif is located in the Scrib LRR region, a domain critical for localization and function, we also added an N-terminal myristoylation signal to negate potential complications due to LRR disruption (Zeitler et al., 2004). The resulting protein, myr-Scrib^TAAA/RAGA^ localized to the basolateral membrane in the follicle epithelium and was enriched at the cell cortex (PM Index > 1), although myr-Scrib^TAAA/RAGA^ localized less well to the cortex than WT myr-Scrib (**Fig. S2B-D**). When expressed in *scrib* mutant wing imaginal discs, myr-Scrib^TAAA/RAGA^ partially rescued the epithelial architecture defects in *scrib* mutants, although growth control was not restored (**Fig. S2E-H**). In follicle cells, myr-Scrib^TAAA/RAGA^ was able to partially rescue the polarity loss phenotype. We observed largely normal apical aPKC localization, with incomplete rescue of epithelial multilayering compared to WT myr-Scrib (**Fig. S2I-J**). These results suggest myr-Scrib^TAAA/RAGA^ retains significant function, and thus that these PP1-interacting consensus motifs are not essential for Scrib’s role in polarity.

### No evidence for physical interaction between Scrib, Dlg and PP1

Given the conserved PP1-interaction motifs in both Scrib and Dlg, we tested whether a physical interaction occurs, first using *in vivo* co-immunoprecipitation (co-IP) assays with transgenic proteins overexpressed in follicle cells. In this assay, we could detect copurification of transgenic Sds22, a known PP1 binding partner (**Fig. 2A**) (Ceulemans et al., 2002). However, Scrib or Dlg were not detected copurifying with PP1 (**Fig. 2A**). We were also unable to reliably detect interaction between Scrib or Dlg and PP1 when combinations of these proteins were overexpressed in cultured *Drosophila* S2 cells, even when cells were crosslinked prior to lysis to stabilize weak and transient protein-protein interactions (**Fig. 2B**).

**Figure 2.**
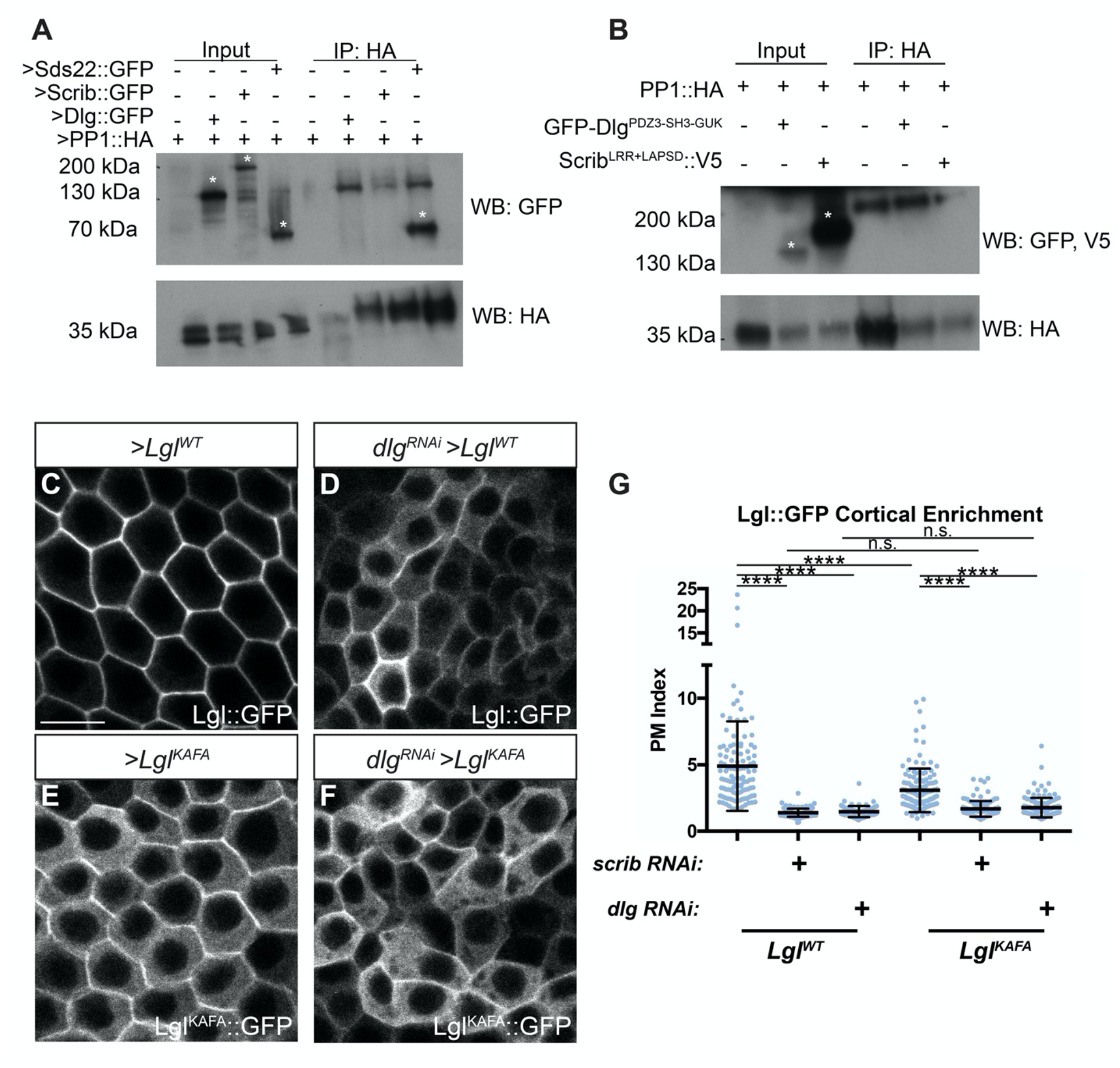
Scrib and Dlg regulate Lgl independently of PP1. (A) CoIP of transgenic Scrib or Dlg and PP1 from follicle cells fails to detect an interaction, although interaction between PP1 and Sds22 is robustly captured. (B) CoIP of overexpressed Dlg or Scrib and PP1 from S2 cells following *in situ* crosslinking also failed to reliably detect interaction between these proteins. Asterisks in (A-B) indicate relevant bands. Lgl^WT^ membrane localization (C) is severely disrupted by *dlg* RNAi (D). Lgl^KAFA^ (E) has increased cytoplasmic localization compared to Lgl^WT^ and is further decreased by *dlg* RNAi (F). (G) Quantification of Lgl membrane localization. Scale bars, 10μm. (G) One-way ANOVA with Tukey’s multiple comparisons test. Error bars represent S.D. PM Index=cortical/cytoplasmic intensity. Data points are measurements from individual cells. n.s. (not significant) P > 0.05, ****P < 0.0001.

### Scrib and Dlg can regulate Lgl independently of PP1

As an additional functional test of the relationship between Lgl regulation by PP1 and its regulation by Scrib and Dlg, we made use of a mutant Lgl protein that cannot interact with PP1 (Lgl^KAFA^) (Moreira et al., 2019). When expressed in the follicle epithelium, Lgl^KAFA^ exhibits an increased cytoplasmic distribution, presumably resulting from its impaired ability to be dephosphorylated and return to the membrane (**Fig. 2C,E,G**) (Moreira et al., 2019). When expressed in *scrib-* or *dlg-*depleted cells rather than WT cells, Lgl^KAFA^ cortical localization was even further reduced, suggesting that even in the absence of PP1 regulation, Lgl^KAFA^ is still dependent on Scrib and Dlg (**Fig. 2F-G**). Interestingly, there was no difference between Lgl^KAFA^ and Lgl^WT^ cortical levels in *scrib-* or *dlg*-depleted cells (**Fig. 2G**). Furthermore, overexpression of PP1 in *scrib-* or *dlg-*depleted cells did not rescue Lgl mislocalization (**Fig. S1F-H**). Together, these data suggest that Scrib and Dlg’s polarity functions include a PP1-independent component.

### A cell culture assay for Scrib recruitment

Although we did not find evidence to functionally implicate PP1 in Scrib/Dlg activity, mutating the Dlg RVxF motif nevertheless caused severe loss of function. We therefore tested other, PP1-independent, functions of this protein region. In addition to protecting Lgl, Dlg also stabilizes Scrib at the cell cortex (Khoury and Bilder, 2020; Ventura et al., 2020). We sought to precisely define the regions of Dlg required to recruit Scrib. To this end, we adapted a previously described induced polarity assay using cultured *Drosophila* S2 cells (Johnston, 2020; Johnston et al., 2009). In this method, transgenic expression of the homotypic cell adhesion protein Echinoid (Ed) is used to create a polarized cortical domain at the contact point between two clustered Ed-expressing cells. By fusing a protein of interest to the Ed intracellular domain, one can create polarized localization of any construct. Importantly, S2 cells do not exhibit native cell-cell adhesion or polarity, although they express a subset of polarity proteins (including Scrib and Dlg) at low to moderate levels. We reasoned that fusing Dlg to Ed would create a discrete cortical domain of polarized Dlg that could recruit endogenous Scrib (**Fig. 3A**). Indeed, an Ed-fused fragment of Dlg encompassing its PDZ3-SH3-HOOK-GUK domains was able to robustly recruit Scrib to the polarity site, compared to a control construct containing Ed alone (**Fig. 3B-D**). Because the same Dlg fragment can provide polarity function *in vivo* (Hough et al., 1997; Lu et al., 2021), the Ed assay provides a useful platform to dissect regions mediating Scrib recruitment by Dlg.

**Figure 3.**
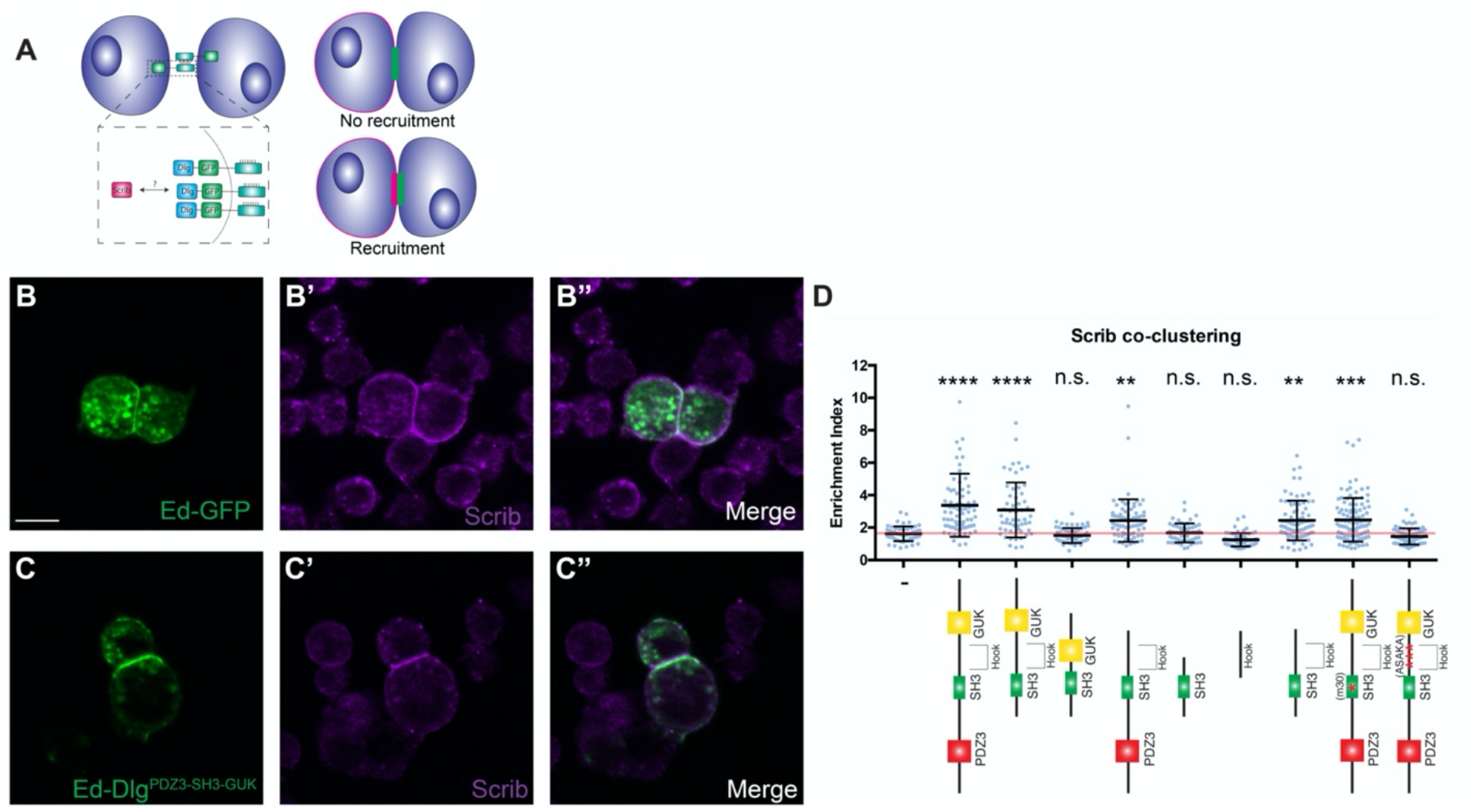
Dlg^SH3-HOOK^ is sufficient for Scrib localization in an induced polarity system. (A) Cartoon of S2 induced polarity assay. Polarizing Dlg by fusion to Ed enables testing of Scrib recruitment in a minimal synthetic system. (B) S2 cells expressing Ed-GFP can be clustered by adhesion between Ed molecules, but this does not alter Scrib localization. (C) When Dlg^PDZ3-SH3-HOOK-GUK^ is fused to Ed, it creates a polarity crescent at the contact site that is able to recruit Scrib. (D) Quantification of Scrib recruitment to the polarity site in various Ed-Dlg constructs schematized below. Dlg^PDZ3-SH3-HOOK-GUK^ is able to enrich Scrib, while the Ed-GFP negative control cannot. The HOOK and SH3 domains are necessary and, when in combination, sufficient to recruit Scrib. Statistical tests are versus the Ed-GFP negative control construct. Red line indicates the average for the Ed-GFP negative control and indicates no Scrib contact site enrichment. Scale bars, 10μm. (D) One-way ANOVA with Tukey’s multiple comparisons test. Error bars indicate S.D. Data points are individual cell clusters. Enrichment index = contact site/non-contact site intensity. n.s. (not significant) P > 0.05, **P < 0.01, ***P < 0.001, ****P < 0.0001.

We tested a series of Ed-Dlg constructs encompassing additional domain truncations and mutations. Consistent with *in vivo* experiments on Dlg function, we found that PDZ3 and GUK were individually dispensable for Scrib recruitment, although GUK deletion resulted in a mild impairment of Scrib recruitment compared to the full-length construct (**Fig. 3D**) (Hough et al., 1997; Khoury and Bilder, 2020; Lu et al., 2021). A construct mimicking the *dlg*^*m30*^ missense mutation in SH3 retained partial ability to recruit Scrib, unlike the *in vivo* situation (Khoury and Bilder, 2020), although it was significantly worse than the WT construct (**Fig. 3D**). We generated a second SH3 domain mutation, designed to disrupt conserved residues that would make up the PxxP binding region of a canonical SH3 domain, and found that this construct also disrupted the ability of Ed-Dlg to recruit Scrib (**Fig. S3A-B**).

### Dlg SH3-HOOK is a minimal fragment necessary and sufficient for polarity *in vivo*

We then turned to the HOOK domain, where the RVxF motif resides. A HOOK-deleted Dlg construct fails to rescue imaginal disc polarity *in vivo*, but this protein localizes to the nucleus rather than the plasma membrane, limiting interpretation (Hough et al., 1997). In the S2 induced polarity assay, the HOOK domain was essential, as a HOOK-deleted construct failed to recruit Scrib (**Fig. 3D**). Interestingly, the same failure was seen with a construct carrying the Dlg^ASAKA^ mutation (**Fig. 3D**). We then tested individual HOOK residues and found that even single amino acid mutations in the RVxF consensus sequence resulted in equivalent disruption of Scrib recruitment activity (**Fig. S3A-B**). In contrast, mutations in evolutionarily conserved residues at the opposite, C-terminal end of the HOOK domain had no effect, suggesting that the HOOK N-terminal region contains the major functional elements (**Fig. S3A-B**). Finally, since single amino acid changes in either the HOOK or SH3 domains disrupt Scrib recruitment, we tested the sufficiency of the domains. Neither domain displayed function alone, but strikingly a fragment composed of SH3-HOOK was sufficient to mediate Scrib clustering (**Fig. 3D**).

We therefore assessed whether this Dlg construct (Dlg^SH3-HOOK^) was also sufficient for function *in vivo*. We compared it to a Dlg protein lacking all three PDZ domains, which Lu et al. 2021 recently demonstrated was sufficient to provide polarity and tumor suppressive activity in *dlg*-deficient follicles and imaginal discs. Our analogous construct (Dlg^SH3-HOOK-GUK^) replicated this result: polarity, architecture, and growth control of discs were also restored in a *dlg* null mutant background (Fig. 4G) (Lu et al., 2021). Excitingly, the smaller Dlg^SH3-HOOK^ also rescued polarity, architecture, and growth control in *dlg-*deficient imaginal discs (**Fig. 4H**). We confirmed this result by taking advantage of a validated *dlg* RNAi line that targets the PDZ2-encoding sequences, allowing us to deplete the endogenous protein but not our transgenes which lack this domain. Depletion of *dlg* in the posterior compartment of wing imaginal discs generates mispolarized tumors, but coexpression of Dlg^SH3-HOOK^ efficiently rescued epithelial polarity, architecture and growth, to a degree indistinguishable from the rescue provided by Dlg^SH3-HOOK-GUK^ (**Fig. S4**). We then tested the constructs in the follicle epithelium. Both transgenic proteins localized to the basolateral membrane, albeit at reduced levels compared to WT Dlg (**Fig. 4A-D**). When expressed in *dlg-*depleted follicle cells (**Fig. 4I**), both Dlg^SH3-HOOK-GUK^ and Dlg^SH3-HOOK^ reduced the basolateral expansion of aPKC to ameliorate polarity and restore monolayer organization (**Fig. 4J-L**), although the former was more efficient than the latter. Intriguingly, even in cases with altered epithelial architecture, Dlg^SH3-HOOK^ restored Scrib recruitment to WT levels, as did Dlg^SH3-HOOK-GUK^ (**Fig. 4M-P**). These data support the conclusion that the SH3 and HOOK domains mediate both Dlg’s Lgl protection and Scrib recruitment activities and that they together constitute a minimal functional unit of the protein that can support epithelial polarity, albeit less efficiently in some tissues than others.

**Figure 4.**
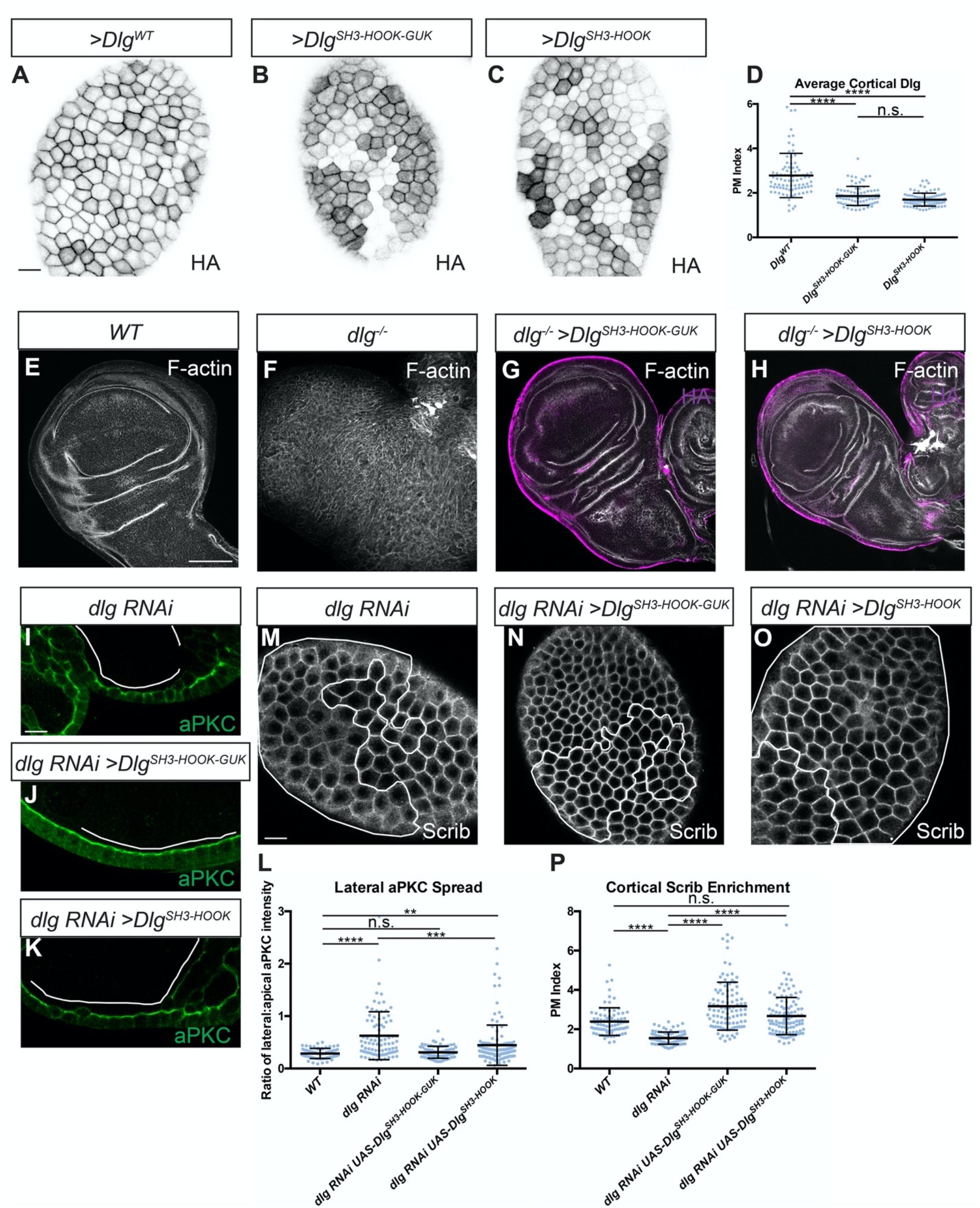
Dlg SH3 and HOOK domains are sufficient for function *in vivo*. (A-C) Like WT Dlg (A), Dlg^SH3-HOOK-GUK^ (B) and Dlg^SH3-HOOK^ (C) localize to the basolateral membrane in follicle cells. All constructs contain an HA epitope tag used for detection. (D) Quantification of cortical localization in (A-C). (E-H) Compared to WT (E) and *dlg* null mutants (F), Dlg^SH3-HOOK-GUK^ (G) and Dlg^SH3-HOOK^ (H) fully rescue polarity and epithelial architecture in wing imaginal discs. (I-K) In monolayered *dlg-*depleted follicle cells (I), both Dlg^SH3-HOOK-GUK^ (J) and Dlg^SH3-HOOK^ (K) provide polarity-rescuing activity, quantitated in (L). Full restoration of monolayering is more efficient by Dlg^SH3-HOOK-GUK^ than Dlg^SH3-HOOK^: 89.5% (n=38) of Dlg^SH3-HOOK-GUK^ show no regions of multilayering in rescued follicles, compared to 20.5% (n=39) of Dlg^SH3-HOOK^ rescued follicles and 0% (n=37) of follicles with *dlg-*depleted clones alone. (M-O). Both Dlg^SH3-HOOK-GUK^ (N) and Dlg^SH3-HOOK^ (O) fully rescue loss of cortical Scrib seen in *dlg*-depleted cells (M), quantified in (P). Scale bars, 10μm except E-H, 100μm. White lines indicate clones of given genotypes. (D, L, P) One-way ANOVA with Tukey’s multiple comparisons test. Error bars indicate S.D. PM Index=cortical/cytoplasmic intensity. aPKC spread is a ratio of lateral:apical fluorescence intensity. Data points are individual cell measurements. n.s. (not significant) P > 0.05, **P < 0.01, ***P < 0.001, ****P < 0.0001.

### The Dlg SH3-HOOK unit regulates Scrib localization, and SH3 provides an additional polarity function

Finally, we investigated the relationship between Dlg’s Scrib recruitment and Lgl protection activities. Nuclear localization of the previous HOOK deletion construct prevented conclusions about its role in the former process. We therefore complemented *dlg* null mutant follicle cells *in vivo* with our HOOK domain missense mutant construct and found that, as in the S2 cell assays, it fails to rescue Scrib cortical localization (**Fig. 5A-D**). The SH3 domain is required for Scrib recruitment *in vivo*, since *dlg*^*m30*^ homozygous cells are defective in recruiting Scrib to the cortex (Khoury and Bilder, 2020). To determine if the SH3 domain is required only for Scrib recruitment, we expressed a membrane-tethered Scrib protein (myr-Scrib) in *dlg*^*m30*^ homozygous follicle cells. Strikingly, this combination yielded a partial rescue of polarity, as assessed by degree of aPKC mislocalization, compared to myr-Scrib in *dlg* null cells (**Fig. 5E-I**). Whereas our previous data show that both SH3 and HOOK domains are required for Scrib localization, this experiment suggests that regions including HOOK cooperate with Scrib to provide Lgl ‘protection’ activity that can be further enhanced by an intact SH3.

**Figure 5.**
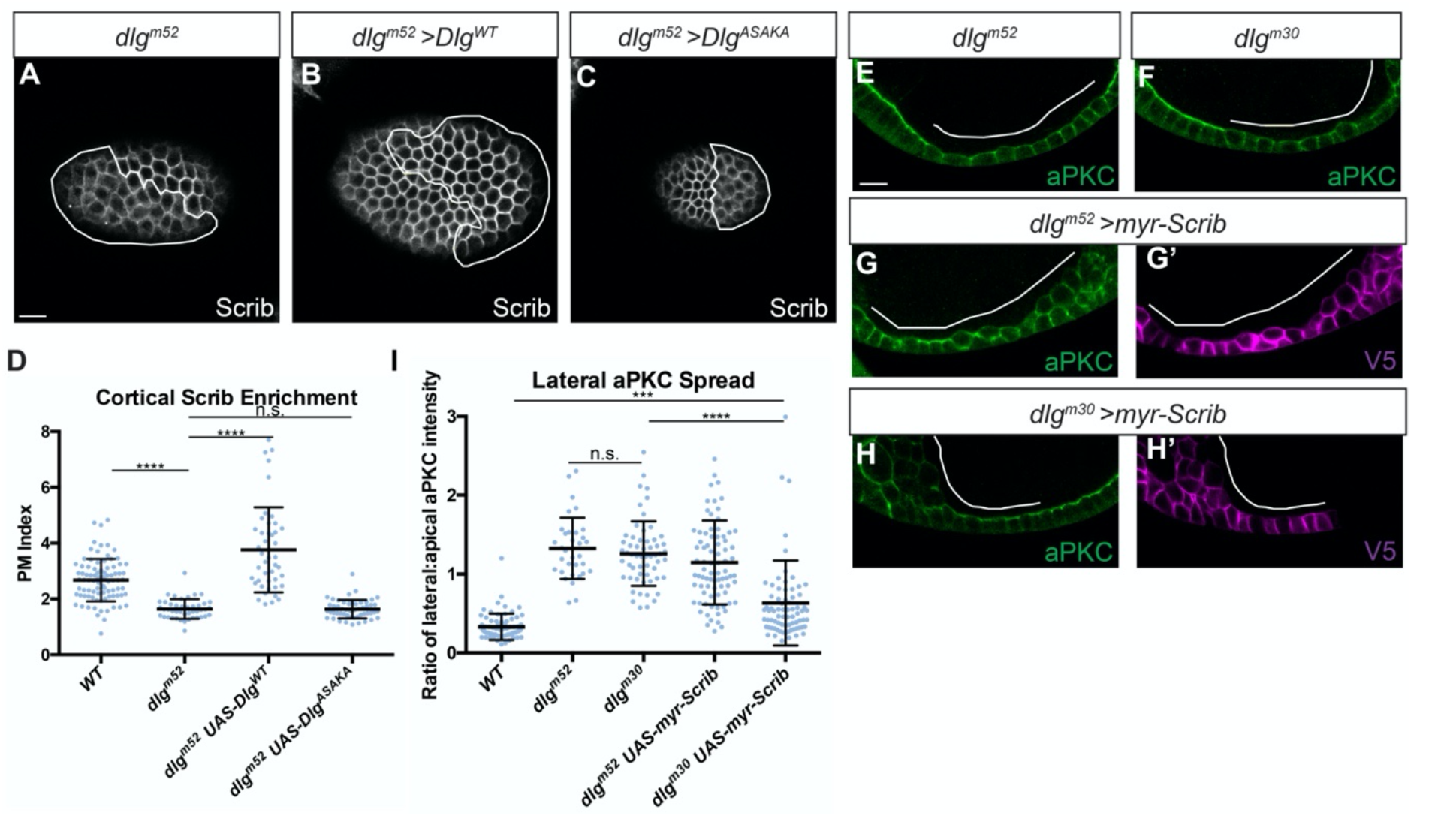
Dlg SH3-HOOK is primarily required to regulate Scrib localization. (A) *dlg*^*m52*^ null mutant cells show reduced cortical localization of Scrib. (B) Scrib mislocalization is rescued by expression of Dlg^WT^. Scrib mislocalization is not rescued by expression of Dlg^ASAKA^ (C). (D) Quantification of Scrib localization in (A-C). (E-H) Both *dlg*^*m52*^ null mutants (E) and *dlg*^*m30*^ SH3 point mutant cells (F) lose polarity and mislocalize aPKC. (H) Preventing Scrib mislocalization by cortical tethering (myr-Scrib) partially suppresses the polarity loss phenotypes of *dlg*^*m30*^ SH3 mutant cells but not *dlg*^*m52*^ null mutant cells (G). (G’-H’) myr-Scrib contains a V5 epitope tag used for detection. (I) Quantification of aPKC mislocalization in (E-H). Scale bars, 10μm. White lines indicate clones of given genotypes. (D,I) One-way ANOVA with Tukey’s multiple comparisons test. Error bars indicate S.D. PM Index=cortical/cytoplasmic intensity. aPKC spread is a ratio of lateral:apical fluorescence intensity. Data points are individual cell measurements. n.s. (not significant) P > 0.05, ***P < 0.001, ****P < 0.0001.

How does SH3-HOOK regulate Scrib recruitment? In cultured mammalian cells it was recently shown that the Scrib LRR and LAPSD domains, which are both necessary and sufficient for polarity function in *Drosophila* (Albertson et al., 2004; Bonello et al., 2019; Choi et al., 2019; Khoury and Bilder, 2020; Zeitler et al., 2004), can co-IP with Dlg1 (Troyanovsky et al., 2021). We tested this Scrib fragment in the S2 induced polarity assay but could not detect recruitment of endogenous Dlg by Ed-Scrib^LRR+LAPSD^ (**Fig. S5A-C**). The ability of Dlg to recruit Scrib in this system, but not vice versa, parallels *in vivo* data showing Scrib localization to be dependent on Dlg, but Dlg localization to be largely independent of Scrib (Khoury and Bilder, 2020; Lu et al., 2021). We were also unable to coIP transgenic Scrib^LRR+LAPSD^ and Dlg^PDZ3-SH3-HOOK-GUK^ from S2 cells (**Fig. S5D**), even with crosslinking and by increasing the starting material used by severalfold. Combined with our induced polarity and *in vivo* genetic experiments, these data support the idea that Dlg recruits Scrib via its SH3-HOOK domains, but that this recruitment may not reflect direct physical binding between the two proteins.

## DISCUSSION

The molecular mechanism of Scrib module function has been a longstanding challenge in the study of cell polarity. Much work has focused on identifying binding partners of Scrib module proteins (reviewed in (Stephens et al., 2018)), with less attention given to the relationships that exist within the Scrib module itself. Here, we perform fine-grained functional analysis of the Dlg protein, defining its minimal required domains. These experiments identified a critical SH3-HOOK module that facilitates Scrib localization and is both necessary and sufficient for Dlg’s polarity activity *in vivo*.

Our search for Scrib module effectors yielded PP1 as an appealing candidate for Lgl regulation. Such a role would be consistent with studies from mammalian cell culture, where both Scrib and Dlg have been found to bind to PP1 (Hendrickx et al., 2009; Nagasaka et al., 2013; Troyanovsky et al., 2021; Van Campenhout et al., 2011; Young et al., 2013), and have been proposed to act as targeting factors that direct PP1 to specific substrates. We failed to find evidence for a physical complex between Scrib, Dlg and PP1 in *Drosophila*, and our data mutating PP1-binding consensus sequences support alternative functions for these motifs, unrelated to PP1-binding. Although we cannot rule out that PP1-Scrib module interactions occur in Drosophila at a low affinity, we note that a direct role for such interactions in regulating mammalian cell polarity remains to be demonstrated. Moreover, our data demonstrate that Scrib and Dlg influence Lgl localization at least partially independently of PP1, which is consistent with the weak phenotype of *pp1* compared to *scrib* module mutants (**Fig. S1A-B**) (Moreira et al., 2019).

Our data using Lgl^KAFA^ are consistent with two possible roles of Scrib and Dlg in polarity. First, Scrib and Dlg could limit Lgl phosphorylation through partners other than PP1, since Lgl mislocalization in *dlg*-depleted cells can be restored by co-depletion of aPKC or by mutating Lgl phosphorylation sites to alanine (Khoury and Bilder, 2020; Ventura et al., 2020). Second, Scrib and Dlg’s regulation of Lgl could involve a phosphorylation-independent component. Consistent with this possibility, we found that a non-phosphorylatable Lgl protein (Lgl^S5A^) still exhibited reduced cortical localization in *scrib-* and *dlg-*depleted cells (**Fig. S6**). These findings draw parallels with the *C. elegans* zygote, where PAR-2 can ‘protect’ PAR-1 both by physical binding as well as competing for aPKC’s activity to reduce PAR-1 phoshorylation (Ramanujam et al., 2018). However, evidence for physical binding of Lgl with Scrib or Dlg in *Drosophila* is currently lacking, outside of a report of binding to the polarity-dispensable GUK domain (Zhu et al., 2014). Thus, although speculative, the analogy of PAR-1 regulation to Lgl ‘protection’ may provide an appealing basis for future experiments. Lastly, although Scrib and Dlg do not require PP1 to regulate Lgl, it is possible that PP1 requires Scrib and Dlg to do so, since the degree of Lgl mislocalization in *scrib-* and *dlg-*depleted cells is not enhanced by removing PP1-dependent regulation (Lgl^KAFA^, **Fig. 2G**).

Dlg is required for Scrib recruitment, proximity assays reliably detect Scrib near Dlg (Nakajima et al., 2019; Sharifkhodaei et al., 2019; Sharp et al., 2021), and an optogenetic relocalization experiment showed that either Scrib or Dlg can induce relocation of the other protein (Ventura et al., 2020). However, we were unable to biochemically detect a Scrib-Dlg complex in extracts from follicles or when the proteins were overexpressed in cell culture. A recent mass spec dataset from *Drosophila* embryos also failed to detect Scrib in Dlg IP samples and vice versa (Nakajima et al., 2019). In flies, biochemical evidence for such a complex involves coIP from synapse-containing tissues such as larval muscle and adult brains (Mathew et al., 2002; Rui et al., 2017); in mammalian cells evidence for coIP comes from cultured cells (Awadia et al., 2019; Troyanovsky et al., 2021). Several of the above cases involve mutual binding partners, and require that partner for coIP, such as Gukholder at the neuronal synapse and SGEF in epithelia (Awadia et al., 2019; Mathew et al., 2002). Given the inconsistent evidence for biochemical interaction, we feel that it is prudent to continue to refer to the Scrib proteins as a ‘module’ rather than a complex.

Our studies identify a critical motif in the Dlg N-terminal HOOK domain that is, in combination with the SH3 domain, required for polarity and Scrib localization. In the wing imaginal disc, SH3 and HOOK are alone sufficient to support full polarity function. In follicle cells, SH3 and HOOK are also sufficient to support Scrib recruitment. Polarity activity in this tissue is less efficient, with full architectural rescue that is less penetrant than with the SH3-HOOK-GUK construct. The SH3-HOOK-rescued follicle cell clones resemble clones mutant for GUK-truncated *dlg* alleles, where polarity in cells retaining epithelial structure is largely normal (Khoury and Bilder, 2020). A follicle-specific role for the GUK domain may involve its known function in spindle orientation, which is required in the follicle but not the wing disc epithelium (Bellaïche et al., 2001; Bergstralh et al., 2013; Bergstralh et al., 2016). Overall, the data demonstrate that SH3 and HOOK domains alone are the minimal elements required for Dlg to recruit cortical Scrib and provide partial polarity function.

How might the SH3 and HOOK domains operate? MAGUK-family SH3 domains are “non-canonical” in that they cannot bind the polyproline ligands bound by typical SH3 domains, because they lack key residues in the PxxP binding pocket (McGee et al., 2001). The HOOK domain is a conserved linker of variable length between the SH3 and GUK domains (Zhang et al., 2013) that is thought to create interdomain allostery, facilitating an intramolecular interaction that enables functions that the individual domains lack in isolation (McCann et al., 2012; McGee and Bredt, 1999; McGee et al., 2001; Rademacher et al., 2019; Zhang et al., 2013). One demonstrated function of HOOK domains is to negatively regulate binding of certain GUK domain ligands, presumably by influencing the SH3-GUK interaction (Golub et al., 2017; Marcette et al., 2009; Qian and Prehoda, 2006). However, the dispensability of the GUK domain for polarity *in vivo* and in the S2 induced polarity assay reveals that such regulation is not important for Scrib recruitment and polarity function. A second function of HOOK is to mediate electrostatic binding to the membrane (Lu et al., 2021), but our experiments mutating non-polar amino acids and supplying membrane tethering in S2 cells show that additional SH3-dependent functions reside in HOOK. Our single amino acid resolution mutant analysis reveals that the HOOK N-terminus is essential to this function, and an appealing model is that it works with the SH3 domain to bind an additional scaffolding factor to permit Scrib recruitment. Once Scrib has been recruited, SH3-HOOK and Scrib are together competent to protect Lgl through an unknown cooperative activity that defines basolateral identity. In support of this model, we find that constitutively tethering Scrib to the membrane can partially bypass a *dlg* SH3 mutant allele, demonstrating that a primary function of SH3 is to recruit Scrib. To our knowledge, this is the first case where a Scrib construct can rescue a *dlg* mutant, providing further evidence for the cooperative nature of Scrib module function in basolateral polarity and pointing to the Dlg SH3-HOOK as a primary mediator of this. Exploring this model will be an important aspect of future studies.

In sum, our in-depth interrogation of the core polarity regulator Dlg defines a minimally sufficient fragment composed of the SH3-HOOK domains, as well as single amino acids in the HOOK domain, that are essential for polarity function. These domains cooperatively recruit Scrib to the cell cortex and supply an additional function that is independent of PP1 that enables Lgl to antagonize aPKC. The data advance our understanding of how basolateral polarity is established and contribute a significant step towards mechanistic understanding of the Scrib module machinery.

## Acknowledgements

We thank E. Morais-de-Sá, C. Johnston and Y. Hong for fly stocks and reagents, L. Mathies for cloning the UAS-Dlg constructs and K. Prehoda, Y. Hong, M. Kitaoka and the Hariharan and Bilder labs for helpful discussions. We thank the UC Berkeley Cell Culture Facility for help with S2 cell experiments. Stocks obtained from the Bloomington Drosophila Stock Center (NIH P40OD018537) and resources from the Drosophila Genomics Resource Center (NIH 2P40OD010949) were used in this study. This work was supported by NIH grant R35 GM130388 to D.B. and AHA Predoctoral Fellowship 20PRE35120150 to M.J.K.

## Competing interests

The authors declare no competing or financial interests.

## MATERIALS AND METHODS

### Fly stocks and genetics

*Drosophila* stocks were raised on cornmeal molasses food at 25**°**C. Mutant alleles and transgenic lines used are listed in **Table S1**. Follicle cell mutant clones were generated using the MARCM technique with hsFLP induction by 37°C heat shock for 1 hour on three consecutive days beginning at 120 hours after egg deposition (AED) for *FRT19A* stocks, and two consecutive days for *FRT82B* stocks. For clonal GAL4 expression, larvae were heat shocked once for 13 minutes at 37°C 120 hours AED to generate flip out clones. For all clonal experiments, newly eclosed females were fed with yeast and dissected three days after eclosion. Unless otherwise noted, pan-follicle cell expression used *traffic jam*-*GAL4* and temperature sensitive *tub-GAL80ts*. After one-two days on yeast, newly eclosed females were shifted to 29°C for three days to induce GAL4 expression before ovary dissection.

### Molecular cloning

To generate *UAS-Dlg*^*ASAKA*^*::HA*, pUASTattB was digested with EcoRI and XbaI and overlapping fragments amplified from the Dlg cDNA were assembled using Gibson assembly. For the *UAS-Dlg*^*SH3-HOOK-GUK*^*::HA* and *UAS-Dlg*^*SH3-HOOK*^*::HA* constructs, pUASTattB was digested with XhoI and XbaI and fragments encompassing the appropriate domains were amplified from the pUASTattB-Dlg^WT^ vector and assembled via Gibson assemnbly. *UAS-myr-Scrib*^*TAAA/RAGA*^*::V5* was generated from *UAS-myr-Scrib::V5* by first using the NEBuilder Gibson Assembly kit to insert a KpnI site 5’ of the myr signal. Then, the resulting plasmid was digested with KpnI and AgeI and fragments containing the desired mutations were amplified and assembled using the NEBuilder Gibson Assembly kit. To generate the Ed-Dlg constructs for S2 cell expression, mutations of interest were introduced into the *pMT-Ed::GFP::Dlg*^*PDZ3-SH3-HOOK-GUK*^ plasmid (Garcia et al., 2014) using the NEBaseChanger site directed mutagenesis kit as directed by the manufacturer (NEB). To generate the cytosolic *pMT-GFP:: Dlg*^*PDZ3-SH3-HOOK-GUK*^ and *pMT-Scrib*^*LRR+LAPSD*^*::V5 p*lasmids, the corresponding regions of the Dlg and Scrib cDNAs were amplified from the *pMT-Ed::GFP::Dlg*^*PDZ3-SH3-HOOK-GUK*^ and *pUASTattB-myr-Scrib::V5* plasmids, respectively. Fragments were then assembled using the NEBuilder HiFi DNA assembly kit (NEB) as instructed into XhoI/EcoRI or AgeI/EcoRI linearized *pMT-His-V5* backbone, respectively. The Dlg sequence used in this study is NP_996405.1 and the Scrib sequence is NP_001036761.3. Primers used for cloning are given in **Table S1**. The deletions made with respect to the Dlg and Scrib reference protein sequences are given in **Table S2**.

### S2 cell culture and induced polarity assay

S2 cells were obtained from the UC Berkeley Cell Culture Facility and cultured using standard methods at 25°C in Schneider’s media (Invitrogen) supplemented with 10% FBS and 1% penicillin/streptomycin. Transfections were performed using the Effectene kit (Qiagen) according to manufacturer’s instructions. 2×10^6^ cells per well of a 6-well plate were transfected with 500ng of DNA per plasmid. Cells were incubated in transfection complexes for 48 hours and then switched into fresh media containing 0.5mM CuSO_4_ to induce expression of the metallothionein promoter for 48 hours before experiments. The induced polarity assay was performed essentially as described previously (Johnston, 2020). After 48 hours of induction, transfected S2 cells were resuspended in 3 mL of fresh media containing 0.5mM CuSO_4._ The cell suspensions were agitated in an orbital shaker at 150 RPM in 6-well plates for 2 hours to induce cell clusters. 1 mL per condition of the clustered cell suspension was then allowed to settle on poly-D-lysine coated coverslips and adhere for 30 minutes. The cells were then fixed for 20 minutes in 4% PFA and processed for immunofluorescence as described below.

### Immunofluorescence and microscopy

Ovaries were dissected in PBS and individual ovarioles were separated prior to fixation in 4% PFA for 20 minutes. Wing imaginal discs were dissected from wandering L3 larvae in PBS and fixed for 20 minutes in 4% PFA. Samples were blocked for 30 minutes to 1 hour in 0.1% PBS-T containing 4% NGS and 1% BSA before staining with primary antibodies overnight at 4°C in blocking buffer. Following 3 washes in PBS-T, samples were incubated in 1:400 fluorescent secondary antibodies (Invitrogen) for 2 hours at room temperature. Primary antibodies used are given in **Table S1**. Imaging was performed on either a Zeiss LSM700 inverted point scanning confocal microscope or an upright Zeiss Axio Imager 2 microscope with Apotome 2 using Plan Apochromat 20x/NA 0.8 or LD C-Apochromat 40x/NA 1.1 W objectives. Uncropped confocal images were 1024×1024 pixels with 2 line averages, and widefield images were 512×512 pixels.

### Image analysis and quantification

Image processing and quantification was performed using FIJI software (Schindelin et al., 2012). To quantify Scrib, Dlg and Lgl cortical localization, a 1.17μm wide rectangular ROI spanning a single cell-cell boundary and a second identical width ROI were measured in en face sections. The ratio of membrane:cytoplasmic fluorescence intensity was computed to define the Plasma Membrane Index (PM Index) (Lu et al., 2021). To quantify aPKC localization, lines along the apical and basolateral membranes were measured in FIJI and the ratio of basolateral:apical intensity was computed to give a measure of lateral mislocalization. To quantify enrichment at S2 cell polarity domains, a 0.39μm wide rectangular ROI spanning the contact site between two S2 cells and a second ROI on a non-contacting section of the membrane were measured and the ratio of contact:non-contact fluorescence intensity was computed to give the Enrichment Index. In all cases measurements were taken from single cells, with averages were calculated for each condition. Figures were assembled with Adobe Illustrator.

### Coimmunoprecipitation and Western blotting

Ovary tissue was lysed in ice cold IP buffer (10mM Tris, 150mM NaCl, 0.5mM EDTA, 0.5% NP-40) (Nakajima et al., 2019) by homogenization. S2 cells were resuspended in ice cold IP buffer and lysed for 30 minutes at 4°C by nutation. Lysates were then cleared by centrifugation at 13,400 x g for 20 minutes at 4°C. Following protein concentration determination by BCA assay (ThermoFisher), 200μg of protein per sample was then loaded onto antibody-conjugated Protein G Dynabeads (ThermoFisher) and rotated overnight at 4°C. The following day, antibody-bead complexes were washed 3 times with lysis buffer before eluting the samples by boiling for 10 minutes in 4x loading dye (Bio-Rad) containing 10% β-mercaptoethanol. 60μg ‘input’ samples were also prepared in the same way by boiling in β-mercaptoethanol-containing loading dye. Antibodies used for IP are listed in **Table S1**. To induce crosslinking, cells were washed several times with ice cold sterile PBS to remove traces of culture media. The cells were then incubated in 2mM disuccinimidyl suberate (DSS, ThermoFisher) in PBS for 30 minutes at room temperature. The crosslinking reaction was then quenched by adding Tris to a final concentration of 20mM and incubating for 15 minutes at room temperature.

Western blotting was performed as previously described (de Vreede et al., 2018). Proteins were separated by SDS-PAGE on 7.5% TGX precast gels (Bio-Rad) before being blotted onto methanol-activated 0.45μm PVDF membranes (GE Healthcare) at 300mA for 1 hour. Membranes were then blocked for 1 hour with TBS-T containing 3% BSA before probing with primary antibodies overnight at 4°C. The following day, membranes were washed 3 times in TBS-T before being incubated in 1:2000 secondary antibodies in blocking buffer for 2 hours at room temperature. Following 3 more washes, blots were imaged by ECL chemiluminance (WesternBright) on HyBlot CL autoradiography film (Denville Scientific). Primary antibodies are listed in **Table S1**.

### Multiple sequence alignment

Protein sequence alignments were created with Clustal Omega (Madeira et al., 2019) and visualized with SnapGene Viewer. The Uniprot sequences used for Dlg were: *H. sapiens* Q12959, *M. musculus* Q811D0, *R. norvegicus* Q62696, *D. rerio* Q5PYH6, *X. tropicalis* Q28C55 and *D. melanogaster* P31007. For Scrib, the sequences used were: *H. sapiens* Q14160, *M. musculus* Q80U72, *R. norvegicus* D3ZWS0, *D. rerio* A0A1L1QZF0, *X. tropicalis* XP_031759453.1 and *D. melanogaster* Q7KRY7.

### Statistical analyses

The statistical tests used for each experiment are described in the corresponding figure legends. No data points were excluded. For each experiment ovaries from at least 5 females were examined, with at least 10 ovarioles being analyzed. All plots show individual data points for all measurements used. All experiments were repeated a minimum of two times. Definitions of significance used are: n.s. (not significant) P > 0.05, *P < 0.05, **P < 0.01, ***P < 0.001, ****P < 0.0001. Data were analyzed using Microsoft Excel and GraphPad Prism 6.

### Data availability

The Dlg APEX2 proteomics dataset was previously published (Sharp et al., 2021) and is available from MassIVE proteomics repository and Proteome Exchange using accession numbers MSV000087186 and PXD025378, respectively. All other reagents and data will be made available upon request.

---

**Figure S1.**
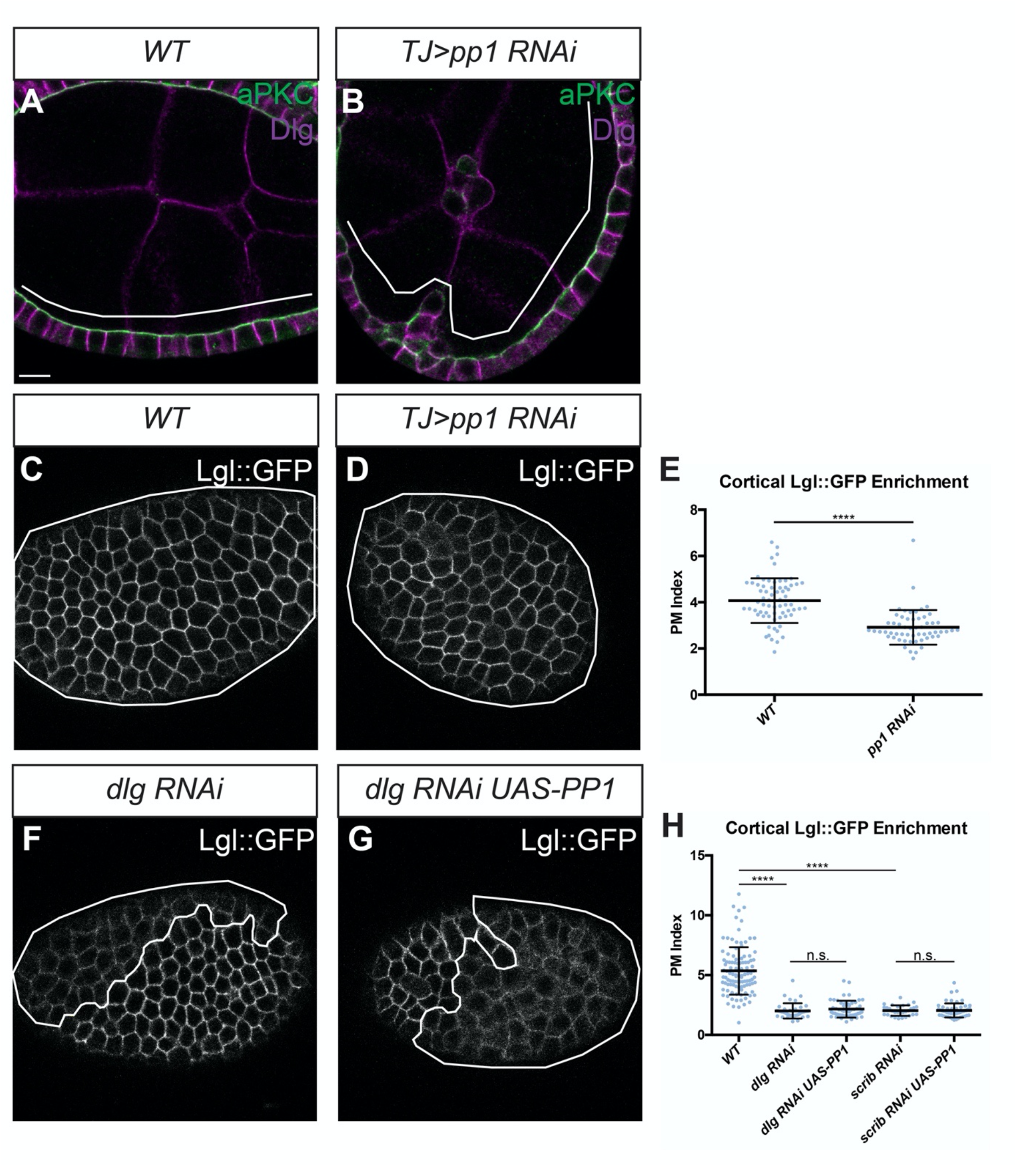
Lgl localization requires both PP1 and Scrib/Dlg. Compared to WT cells (A), *pp1*-depleted cells (B) exhibit mild polarity loss and occasional multilayering. Compared to WT cells (C), *pp1*-depleted cells display mild loss of cortical Lgl (D, also compare to *dlg*-depletion in F). (E) Quantification of Lgl localization. (F) *dlg*-depleted cells strongly mislocalize cortical Lgl and this is not rescued by overexpression of PP1 (G). (H) Quantification of Lgl localization. Scale bars, 10μm. White lines indicate clones of given genotypes. (E) Two-tailed t-test with Welch’s correction. (H) One-way ANOVA with Tukey’s multiple comparisons test. Error bars indicate S.D. PM Index=cortical/cytoplasmic intensity. Data points are individual cell measurements. n.s. (not significant) P > 0.05, ****P < 0.0001.

**Figure S2.**
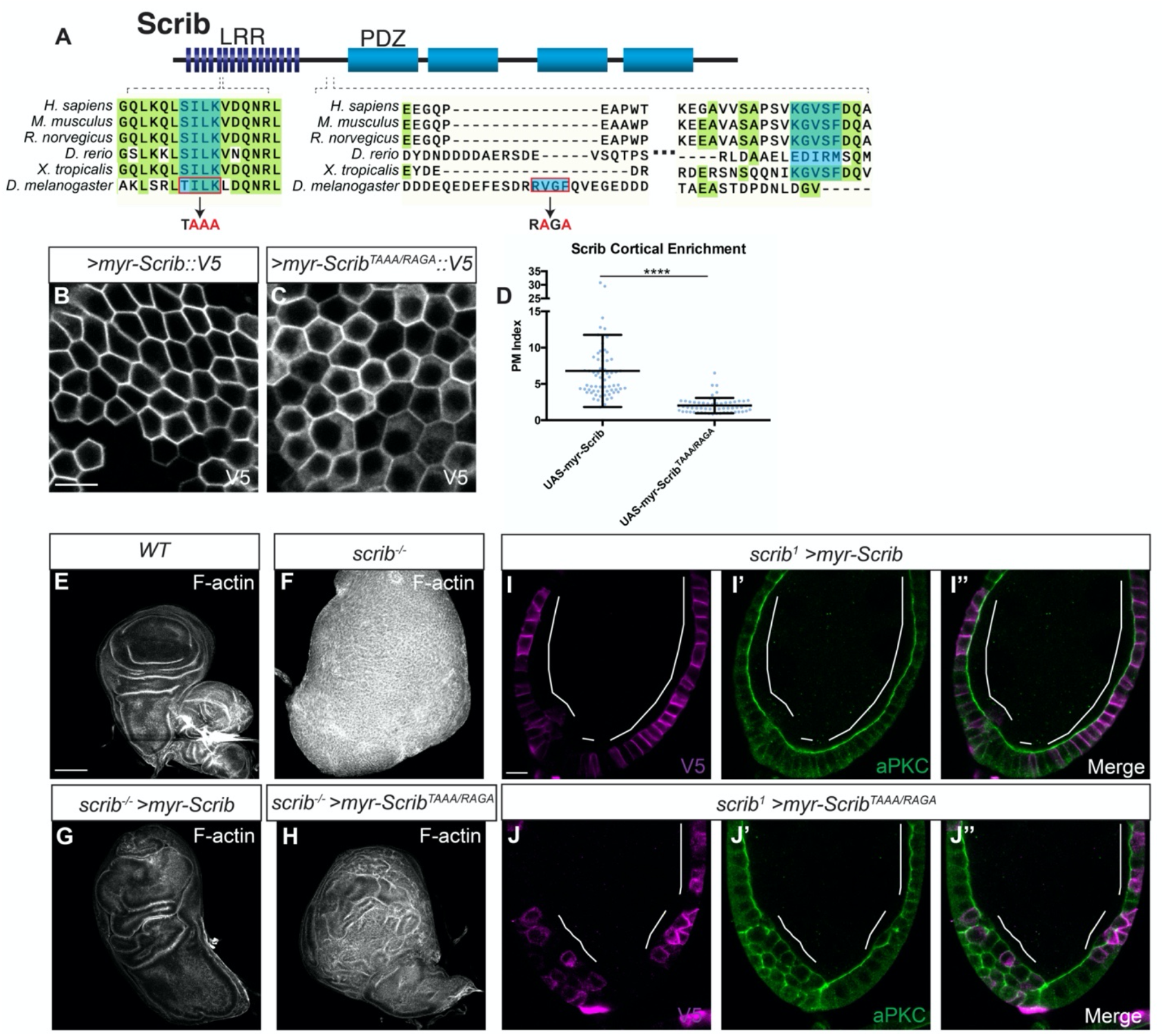
Scrib PP1-binding consensus motifs are partially required for function. (A) Cartoon showing the Scrib protein domain composition and location of the SILK and RVxF motifs. Below: alignment showing conservation of the SILK motif and RVxF motifs. Note that in vertebrates (right), the RVxF motif is located slightly C-terminal to its position in insects (left). Red boxes indicate residues mutated in myr-^ScribTAAA/RAGA^ construct. Compared to WT myr-Scrib (B), myr-Scrib^TAAA/RAGA^ (C) localizes less well to the cell cortex but is still enriched at the basolateral membrane, quantified in (D). Both constructs contain V5 epitope tags, used for detection. (E-H) Compared to WT wing discs (E), *scrib* mutant wing discs overgrow and form tumors (F). Overexpression of myr-Scrib largely rescues these phenotypes (G), while expression of myr-Scrib^TAAA/RAGA^ only partially rescues the *scrib* mutant phenotype (H). (I-J) Compared to myr-Scrib (I), myr-Scrib^TAAA/RAGA^ (J) provides less efficient rescue of *scrib* mutant: myr-Scrib shows complete restoration of the monolayered epithelium in 78.6% (n=14) of follicles, compared to complete restoration in 36.8% (n=19) of myr-Scrib^TAAA/RAGA^ -rescued follicles. Scale bars, 10μm, except E-H, 100μm. White lines indicate clones of given genotypes. (D) Two-tailed t-test with Welch’s correction. Error bars indicate S.D. PM Index=cortical/cytoplasmic intensity. Data points are individual cell measurements. ****P < 0.0001.

**Figure S3.**
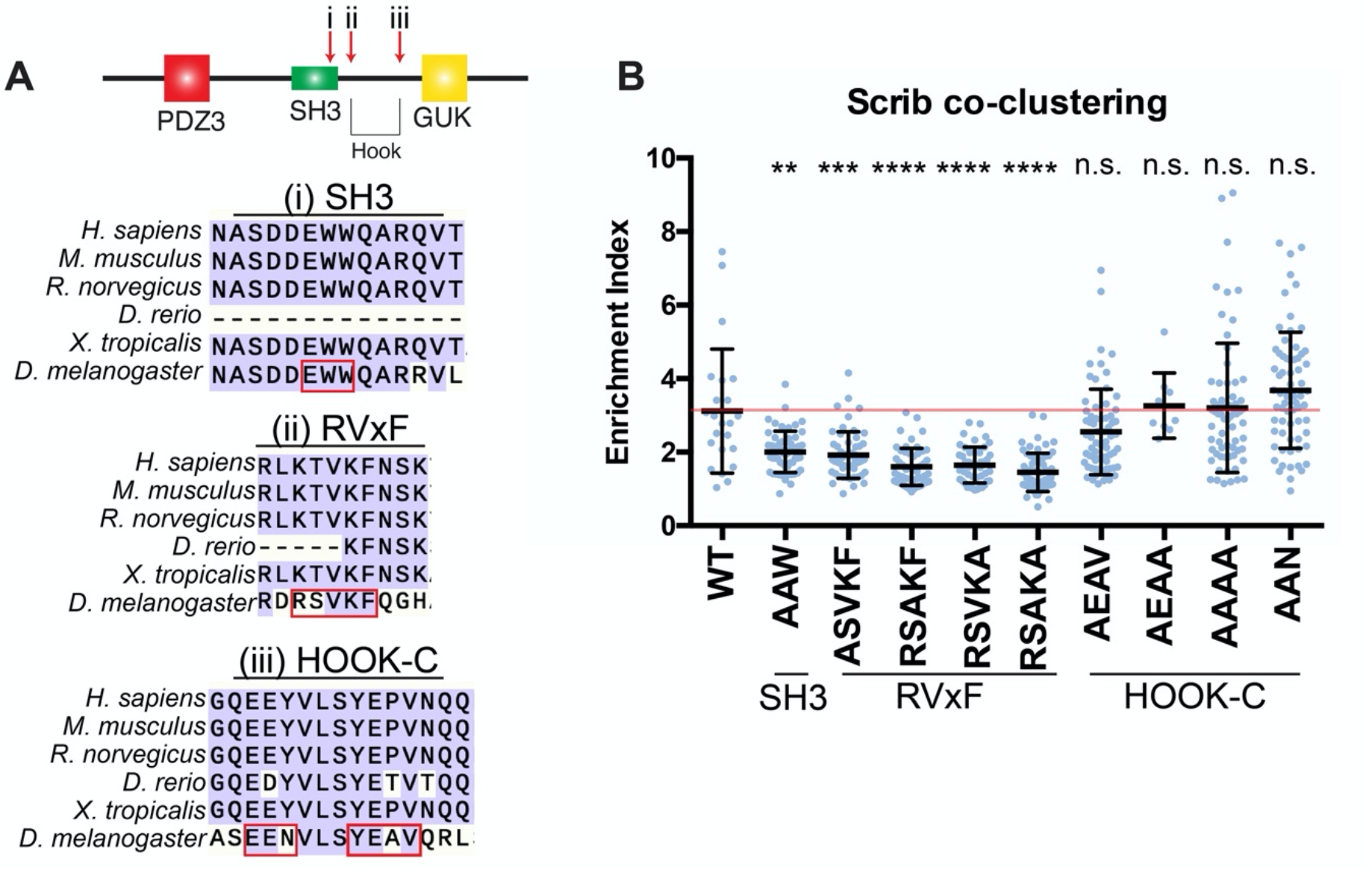
In depth examination of Dlg SH3 and HOOK domains. (A) Schematic of the Dlg domains used in the Ed-Dlg^PDZ3-SH3-HOOK-GUK^ construct, with sequence alignments showing conservation of the SH3 and HOOK domain sequences chosen for study. Motifs targeted for mutation are indicated by red outlines. Arrows in cartoon indicate relative locations of targeted sequences in the protein. (B) Quantification of Scrib recruitment to the polarity site in S2 induced polarity assay. Compared to the WT Ed-Dlg^PDZ3-SH3-HOOK-GUK^ construct, which does recruit Scrib, the SH3 mutant AAW construct has reduced ability to recruit Scrib. Similarly, all four constructs targeting single residues of the RVxF motif show equally impaired ability to recruit Scrib. However, the four constructs targeting conserved residues in the C-terminal HOOK domain do not impair Scrib recruitment. Red line indicates the average for the WT construct control. One-way ANOVA with Tukey’s multiple comparisons test. Error bars indicate S.D. Data points are individual cell clusters. Statistical tests are comparisons to the Ed-Dlg^PDZ3-SH3-HOOK-GUK^ control construct. Enrichment index = contact site/non-contact site intensity. n.s. (not significant) P > 0.05, **P < 0.01, ***P < 0.001, ****P < 0.0001.

**Figure S4.**
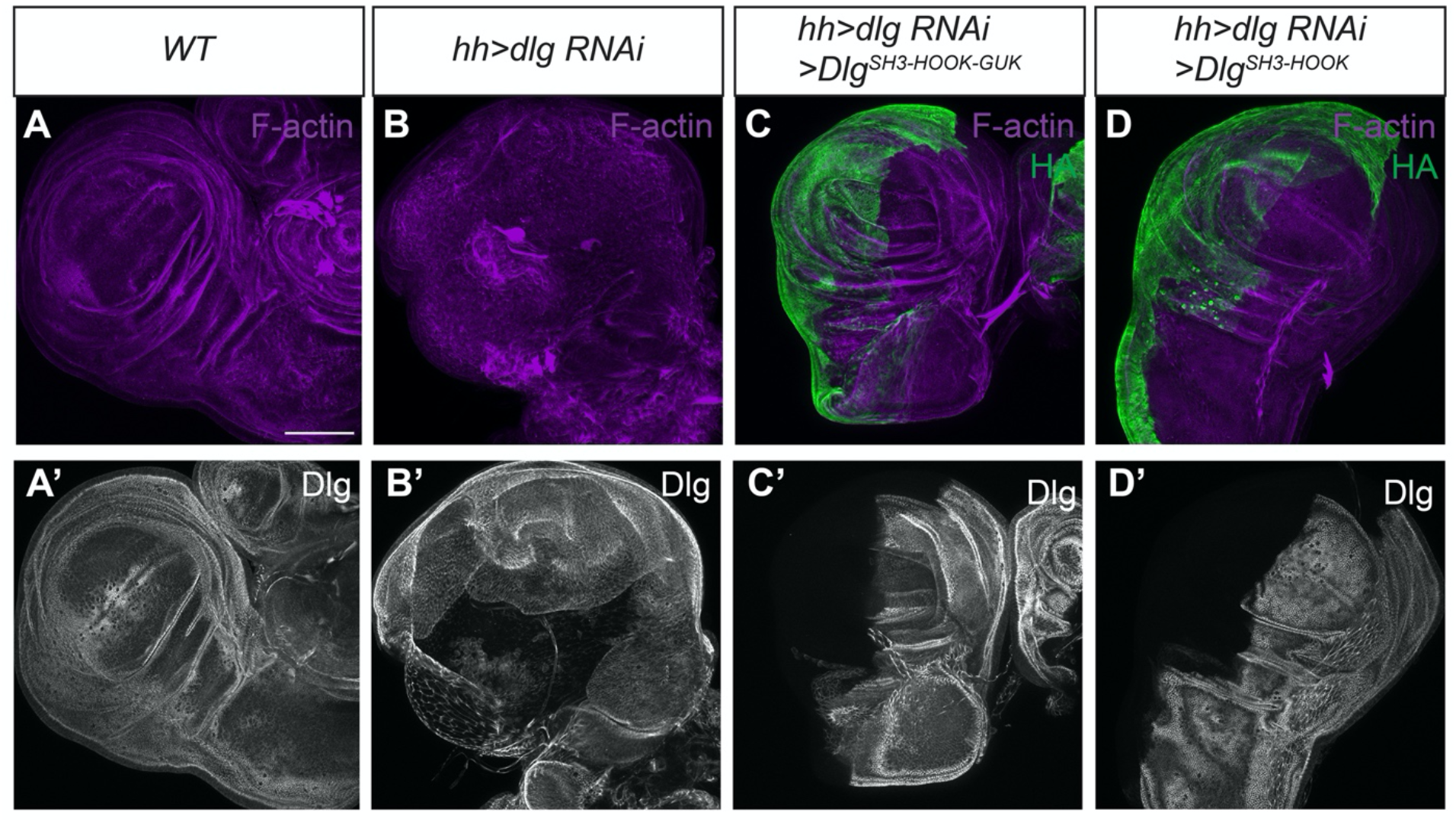
Validation of RNAi rescue approach for minimal Dlg constructs. (A-D) Knocking down *dlg* in the posterior half of the wing disc using an RNAi construct targeting PDZ2-encoding sequences (B) causes polarity loss and disrupted epithelial architecture. These phenotypes are fully rescued by co-expression of Dlg^SH3-HOOK-GUK^ (C) and Dlg^SH3-HOOK^ (D), and neither constructs is targeted by the RNAi reagent used to deplete endogenous Dlg (C’, D’). Scale bars, 100μm.

**Figure S5.**
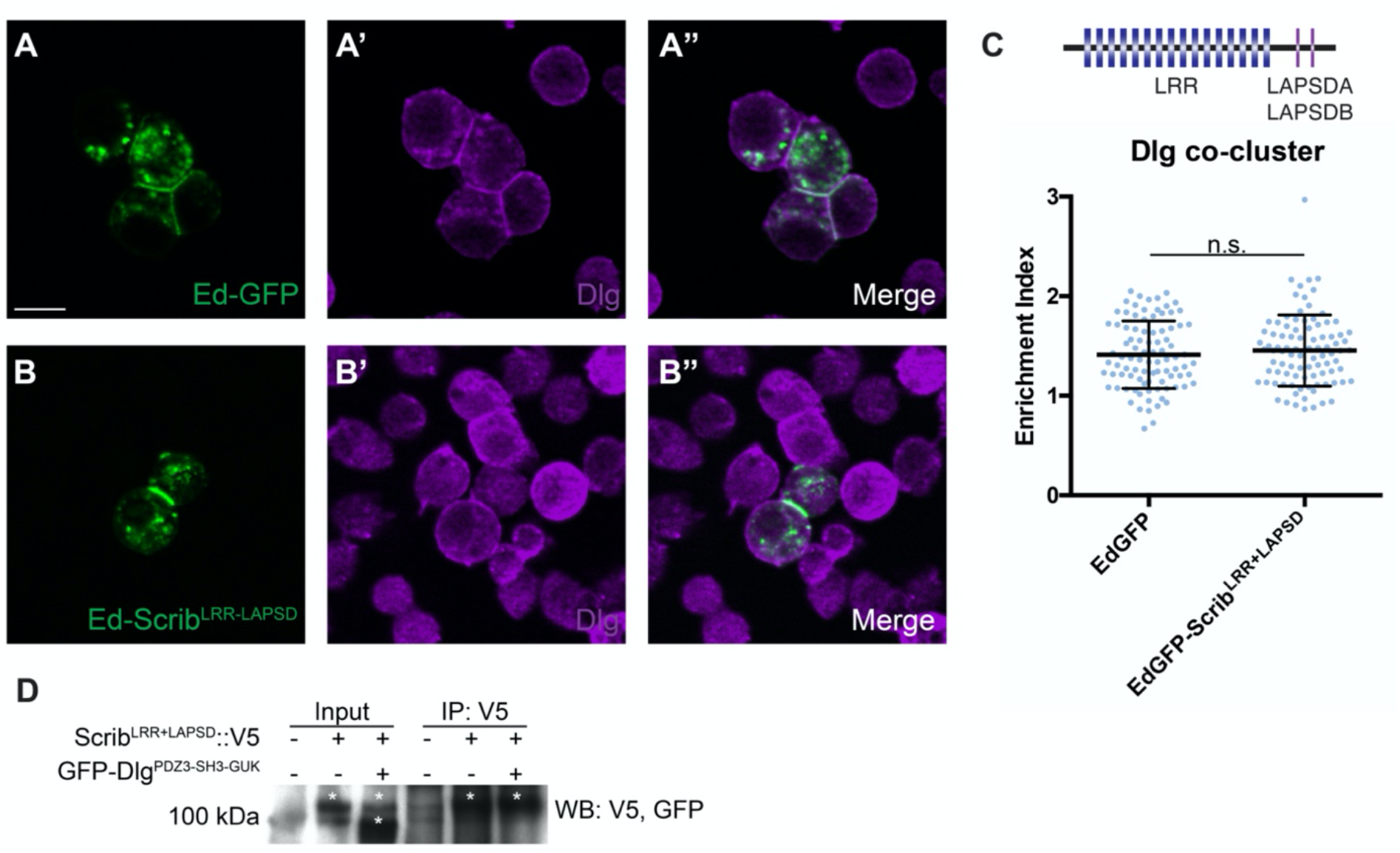
Using S2 cell induced polarity to study Scrib-Dlg interaction. (A) Ed-GFP expression in S2 cells allows induction of a polarity domain where cells adhere. (B) Fusing the Scrib LRR+LAPSD domains to Ed creates a domain of polarized Scrib. (C) Schematic of the Scrib domains used in the Ed-Scrib^LRR+LAPSD^ construct. Quantification of Dlg enrichment shows that Scrib cannot recruit Dlg to the polarity site. (D) CoIP assay of Scrib^LRR+LAPSD^ and Dlg^PDZ3-SH3-HOOK-GUK^ from S2 cells fails to detect interaction between these proteins. Scale bars, 10μm. (C) Two-tailed t-test with Welch’s correction. Error bars indicate S.D. Data points are individual cell clusters. Enrichment index = contact site/non-contact site intensity. n.s. (not significant).

**Figure S6.**
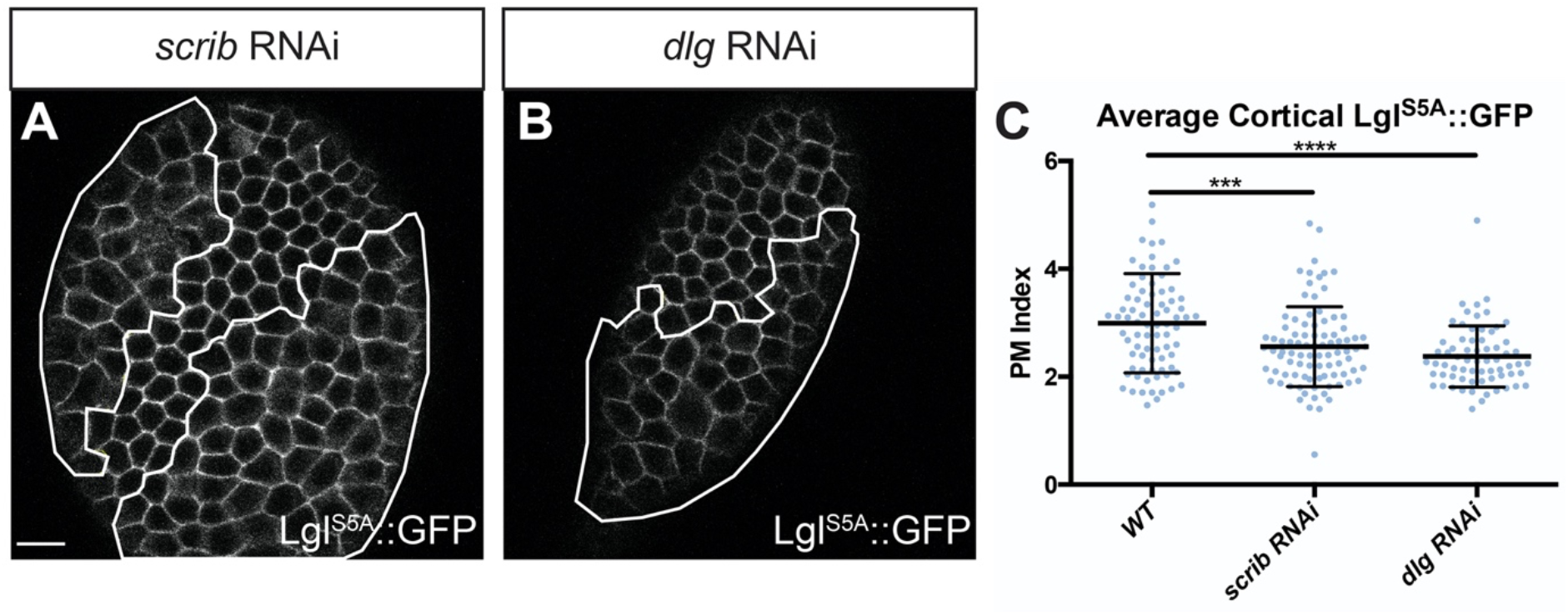
Scrib and Dlg protection of Lgl is partly independent of phosphorylation. (A-B) Compared to its localization in WT cells, non-phosphorylatable Lgl^S5A^ exhibits a slight, but significant, reduction in cortical levels in *scrib* RNAi (A) and *dlg* RNAi expressing cells (B). (C) Quantification of LglS5A::GFP levels. *scrib* or *dlg* RNAi both significantly reduce Lgl^S5A^ cortical levels compared to WT cells. Scale bars, 10μm. White lines indicate clones of given genotypes. PM Index=cortical/cytoplasmic intensity. Data points represent individual cell measurements. Error bars represent S.D. (C) One-way ANOVA with Dunnett’s multiple comparisons test. ***P < 0.001, ****P < 0.0001.

**Table S1.**
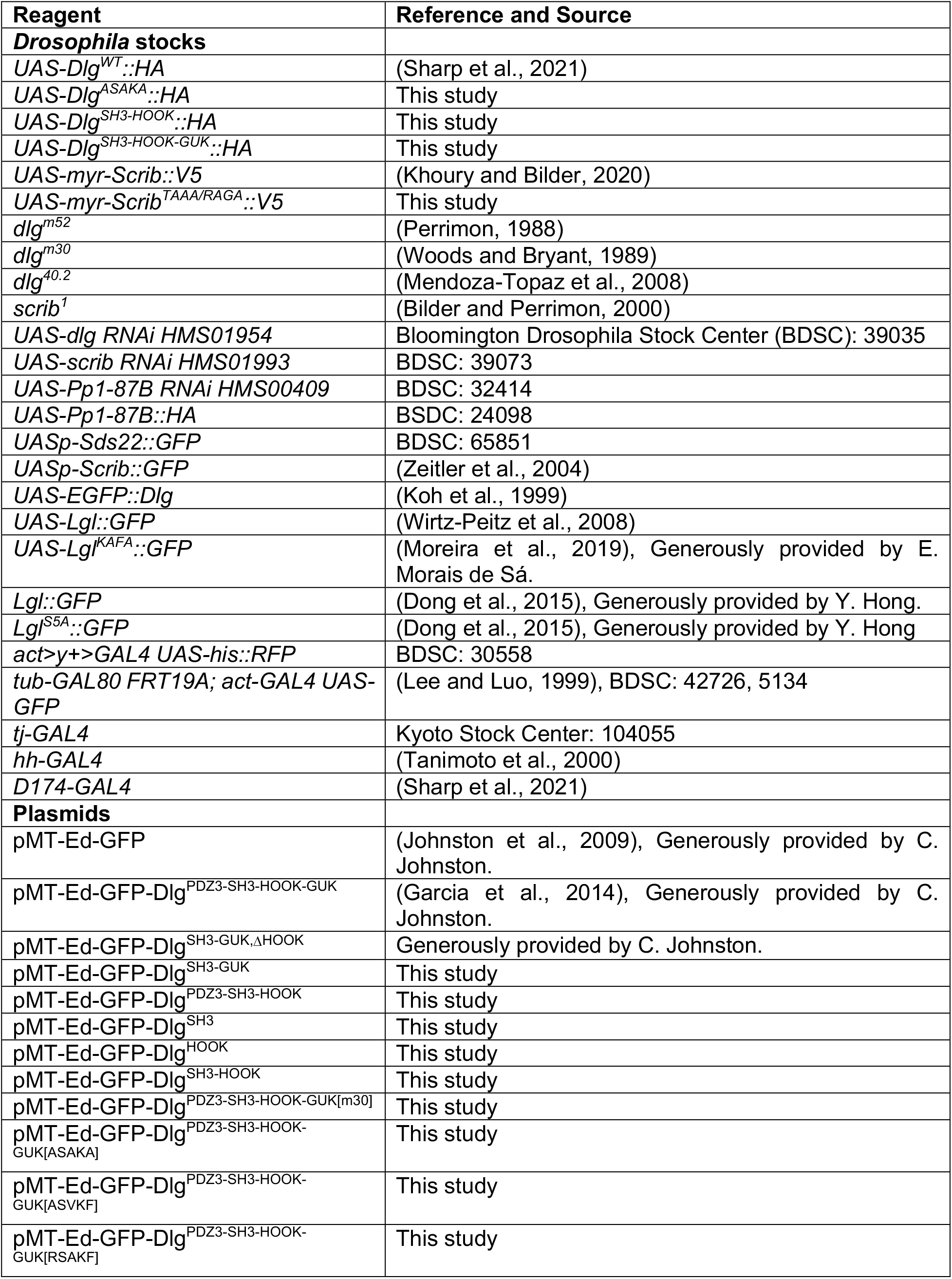
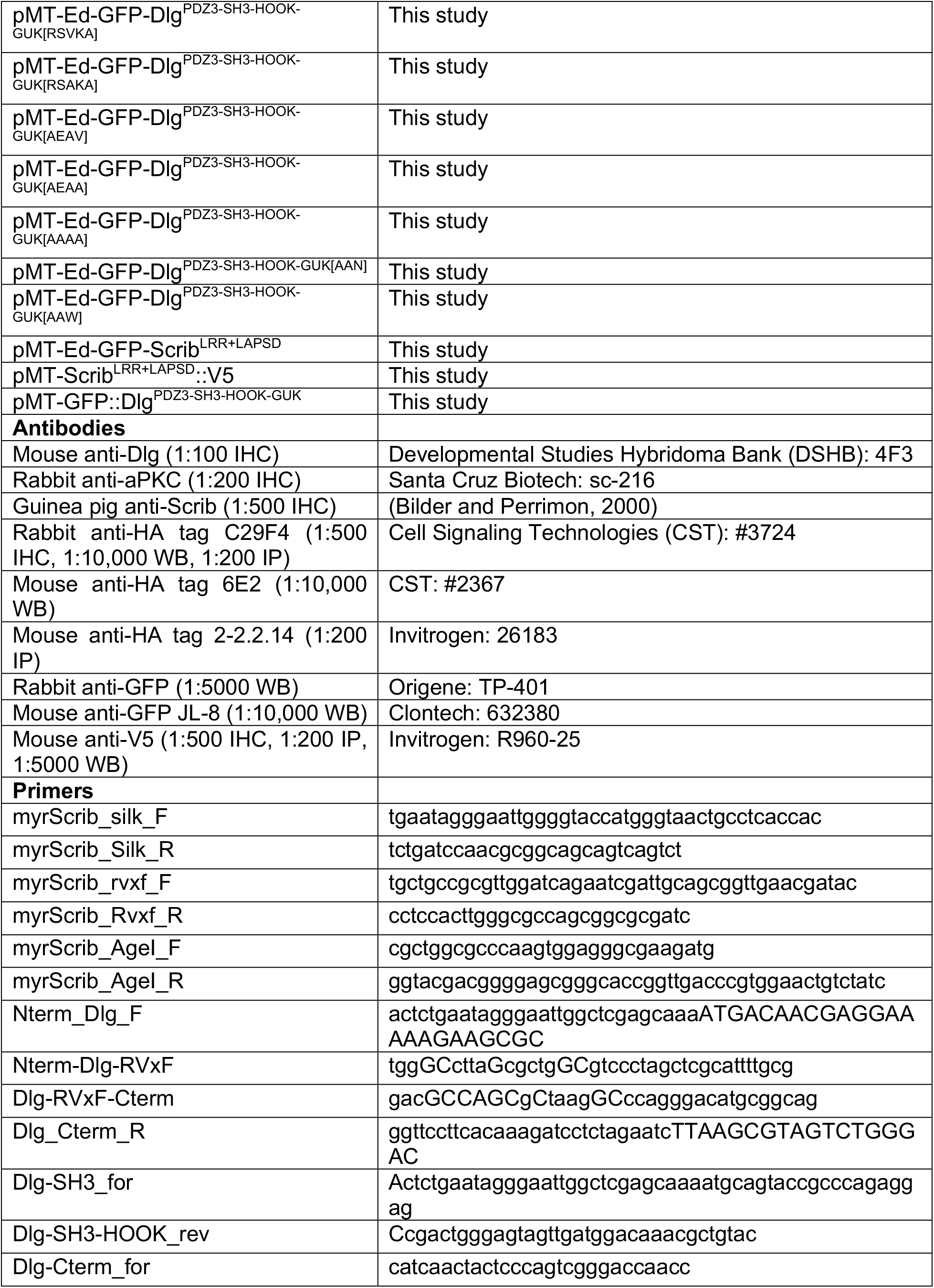
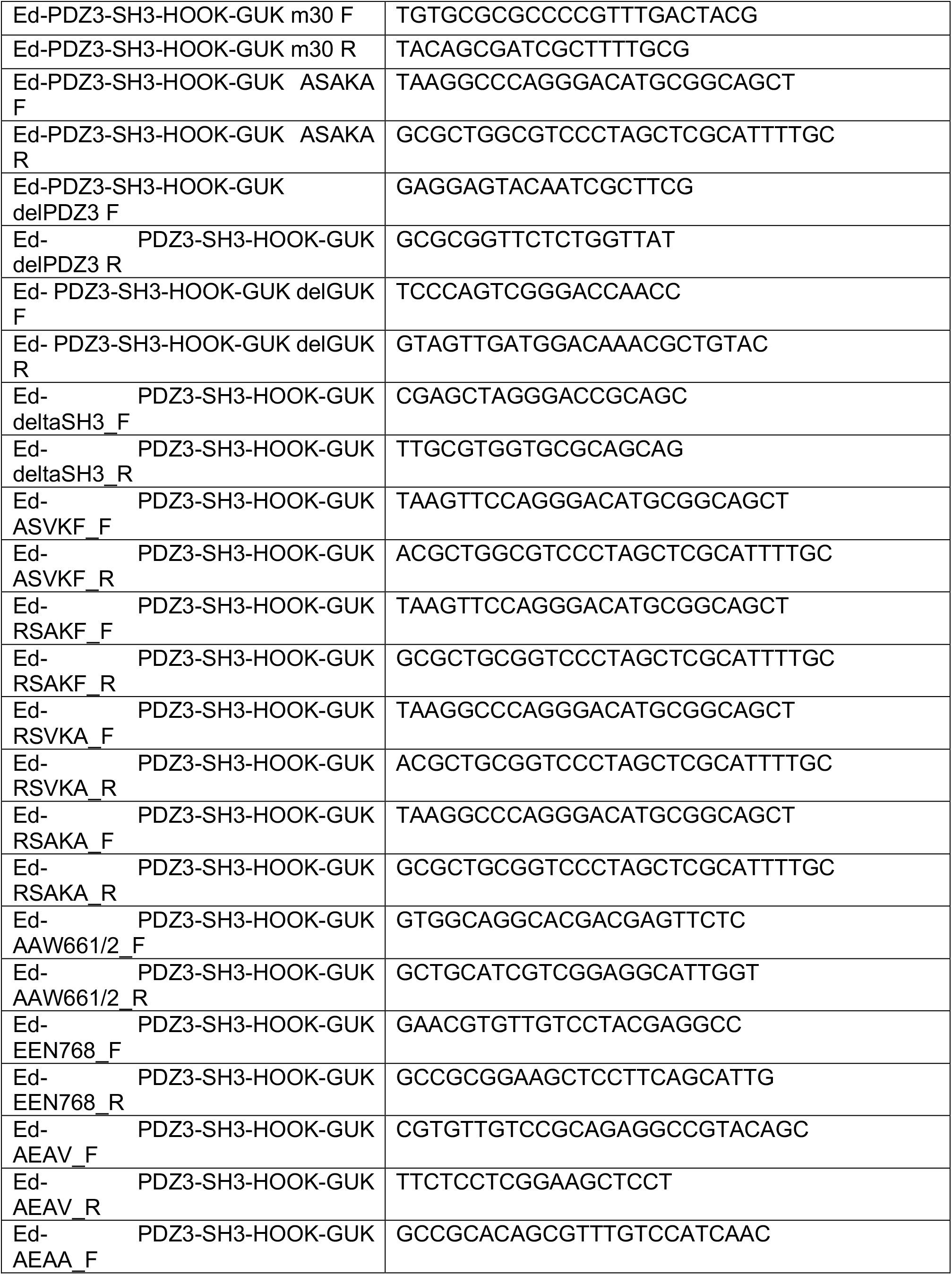
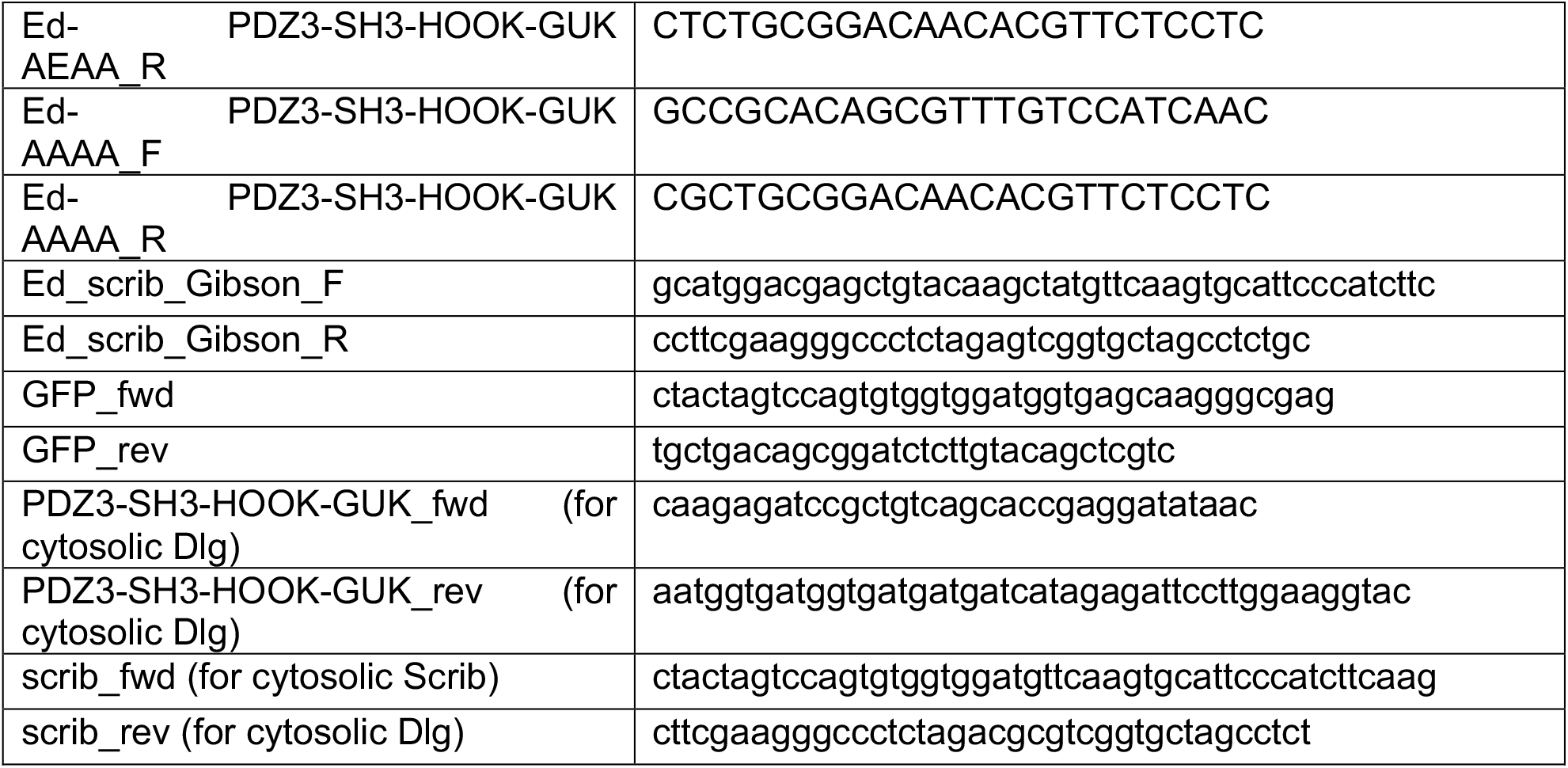
Key Resources.

**Table S2.**
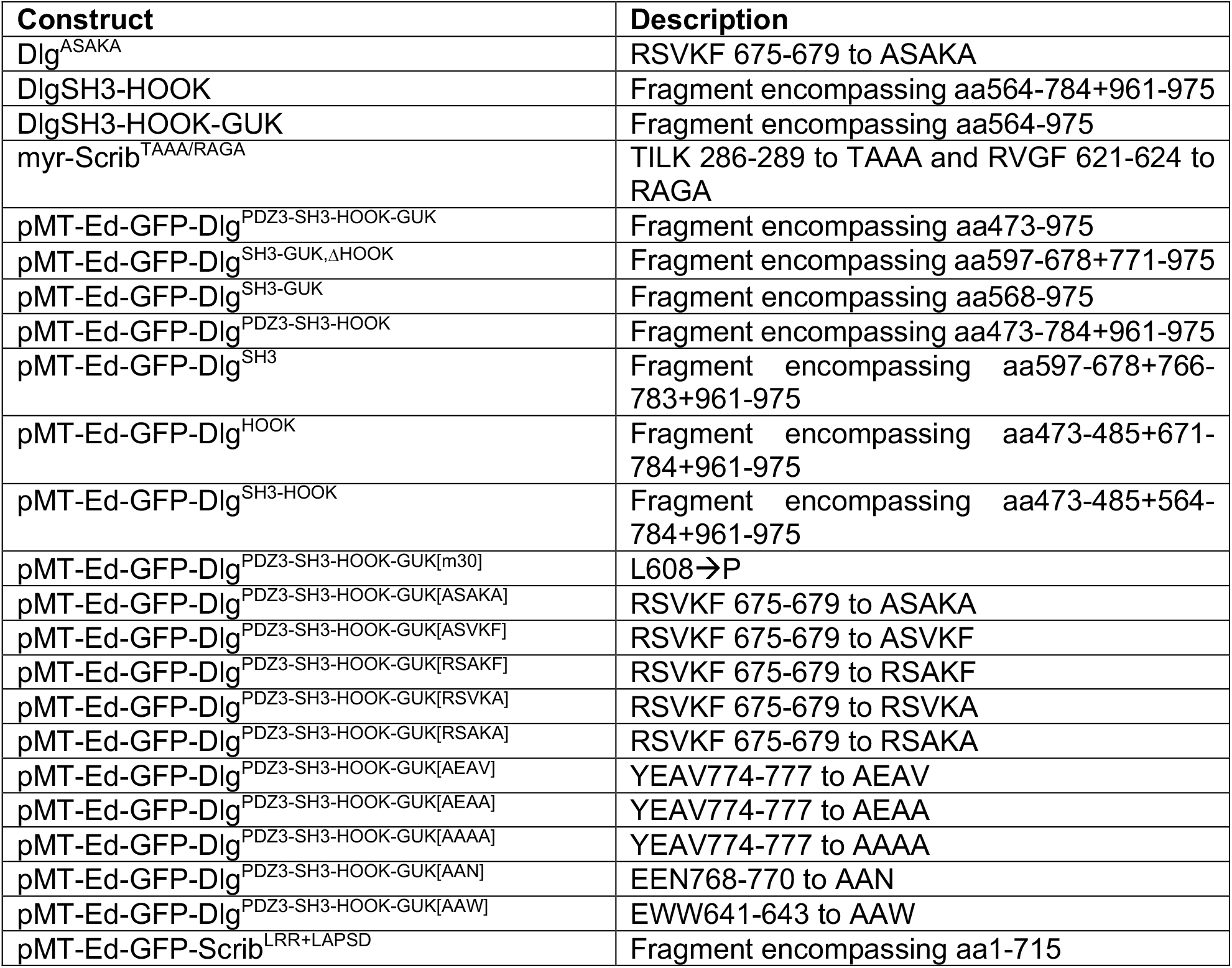
Scrib and Dlg transgenic constructs.

## Notes

### Competing Interest Statement

The authors have declared no competing interest.

